# Transient nuclear deformation primes epigenetic state and promotes cell reprogramming

**DOI:** 10.1101/2021.05.19.444886

**Authors:** Yang Song, Jennifer Soto, Binru Chen, Weikang Zhao, Tyler Hoffman, Ninghao Zhu, Qin Peng, Chau Ly, Pak Kin Wong, Yingxiao Wang, Amy C. Rowat, Siavash K Kurdistani, Song Li

**Affiliations:** Department of Bioengineering, University of California Los Angeles, Los Angeles, CA, 90095, USA; Department of Biomedical Engineering, The Pennsylvania State University, University Park, PA 16802, USA; Department of Bioengineering, University of California San Diego, San Diego, CA, 92093, USA; Department of Integrative Biology & Physiology, University of California Los Angeles, Los Angeles, CA, 90095, USA; Department of Biological Chemistry, David Geffen School of Medicine, University of California Los Angeles, Los Angeles, CA, 90095, USA; Department of Medicine, University of California Los Angeles, Los Angeles, CA, 90095, USA

**Keywords:** Direct cell conversion, histone methylation, DNA methylation, epigenetic regulation, mechanobiology

## Abstract

Cell reprogramming has wide applications in tissue regeneration, disease modeling and personalized medicine, but low reprogramming efficiency remains a challenge. In addition to biochemical cues, biophysical factors can modulate the epigenetic state and a variety of cell functions. However, how biophysical factors help overcome the epigenetic barrier for cell reprogramming are not well understood. Here we utilized microfluidic channels to induce a transient deformation of the cell nucleus, which caused the disassembly of the nuclear lamina and a downregulation of DNA methylation and histone (H3K9) for 12-24 hours. These global decreases of heterochromatin marks at the early stage of cell reprogramming strikingly enhanced the conversion of fibroblasts into neurons and induced pluripotent stem cells. Consistently, inhibition of DNA methylation and H3K9 methylation partially mimicked the effects of mechanical squeezing on iN reprogramming efficiency. Knocking down lamin A had similar effects to squeezing on enhancing the reprogramming efficiency. Based on these findings, we developed a scalable microfluidic system that enabled a continuous cell processing to effectively prime the epigenetic state for cell reprogramming, demonstrating the potential of mechano-biotechnology for cell engineering.

## Introduction

Cell reprogramming technologies, such as somatic cell nuclear transfer, induced pluripotent stem cell (iPSC) reprogramming, and direct reprogramming, can be used to derive desirable cell types that have wide applications in regenerative medicine, disease modeling and drug screening^1–3^. Direct reprogramming not only facilitates the conversion of one cell type into distantly related cell type(s), but also provides a faster and more direct method to obtain a desired cell type by circumventing the pluripotent stage and time-consuming differentiation process. Indeed, fibroblasts can be directly converted into other cell types such as neurons^4^, cardiomyocytes^5^, β-islet cells^6^, blood cell progenitors^7^, and hepatocytes^8^. However, the low yield of reprogrammed cells has limited the translation of direct reprogramming into clinical settings and pharmaceutical applications. For example, mouse neonatal fibroblasts can be converted into induced neuronal (iN) cells via ectopic expression of three transcription factors: Ascl1, Brn2 and Myt1l (BAM), with an efficiency of 1.8-7.7%, but with a much lower efficiency for adult fibroblasts^9^.

A critical step in cell reprogramming is to overcome the epigenetic barrier of heterochromatin and turn on the endogenous genes for cell type conversion. Most of the previous studies have focused on the roles of transcriptional factors and biochemical factors in cell reprogramming^10,11^, but the effects of biophysical factors are much less understood. Cells experience mechanical stimuli at both short and long time scales, from seconds to days, which may result in mechano-chemical signaling, cytoskeleton reorganization and chromatin changes^12–17^. For example, surface topography induces an elongated nucleus shape and increases histone H3 acetylation (AcH3) and H3K4 methylation during fibroblast reprogramming into iPSCs^18^, and three-dimensional (3D) collagen gel increases H3K4 methylation in T cells^19^. In addition, soft matrix decreases H3K9me3 in tumor cells in a cell type-dependent manner^20^, and persistent uniaxial stretching of adhesive substrates decreases H3K9me3 in epidermal cells^21^. Interestingly, compression on the side of adherent mesenchymal cells enhances histone acetylation, while compression on the top of adherent fibroblasts leads to an increase of heterochromatin^22,23^. These differential responses to various biophysical cues suggest that mechanotransduction to the nucleus is context-dependent in adherent cells, which may be attributed to the differences in specific biophysical cues, cell types, and cell adhesions and cytoskeleton organization. Regardless of all these variations, we postulated that an appropriate mechanical perturbation of cell nucleus could induce chromatin remodeling and help overcome the heterochromatin barrier for cell reprogramming. Cells in suspension offer a valuable model to test this hypothesis, in which the complexity of extracellular signals (e.g., adhesion polarity, matrix stiffness, ligand presentation) and intracellular components (focal adhesion complex, cytoskeleton organization) are removed or reduced. Therefore, to directly determine the effect of nuclear deformation on chromatin remodeling, we investigated whether mechanically squeezing suspended cells could regulate the epigenetic state and cell reprogramming. Microfluidic devices have been widely used to study cancer cell responses to cell deformation in confined space^24,25^. Here we developed scalable microfluidic devices to induce nuclear deformation and chromatin remodeling, and explored the translation of the findings into mechano-biotechnology applications.

## Results

To investigate the effect of nuclear deformation on direct reprogramming, we developed a microfluidic device with constriction microchannels, and forced cells in suspension to flow through these channels (**Fig. 1a** and **Supplementary Fig. S1**). To determine a proper channel dimension that would enable nuclear deformation while maintaining cell viability, we considered the size of mouse fibroblasts (19.7 ± 4.5 μm) and their nuclei (10.5 ± 1.2 μm) (**Fig. 1b** and **c**), and fabricated microdevices that had parallel constriction channels with the same height (15 μm) but different widths (3, 5, 7 and 9 μm). Cells passing through wide microfluidic channels (200 μm) did not show significant nuclear deformation (**Supplementary Fig. S2**), and were used as a control in all studies. To determine the effect of nuclear deformation on cell membrane integrity, a fluorescently-tagged antibody (Cy5-IgG) was added to the culture media of fibroblasts in suspension before the cells were introduced into the microfluidic device. As shown in **Fig. 1d**, Cy5-IgG could be detected in many cells that passed through 3 µm- and 5 µm-wide channels, indicating a cell membrane rupture^24^, which was not detected in cells passing through 7 μm-wide channels. Additionally, by using fibroblasts expressing histone 2B (H2B)-GFP, we observed that 3 µm and 5 µm-wide channels, but not 7 μm-wide channels, induced nuclear blebbing or disrupted the nuclear lamina after squeezing, as shown by histone leakage from the nucleus (**Fig. 1d**). To further assess how channel width affected the cell viability, we performed live/dead cell staining and PrestoBlue assays. In comparison to the control group (passing through 200-μm channel), 7 and 9 μm-wide channels did not induce noticeable cell death at 3 hours and one day after mechanical deformation **(Fig. 1e** and **Fig. S3-S4)**. In contrast, after passage through 3 µm- and 5 µm-wide channels, there was a significant decrease in cell viability. These results suggested that 3 µm- and 5 µm-wide channels induced cell damage. Therefore, 7-μm channels were used in the follow-up studies.

**Fig. 1.**
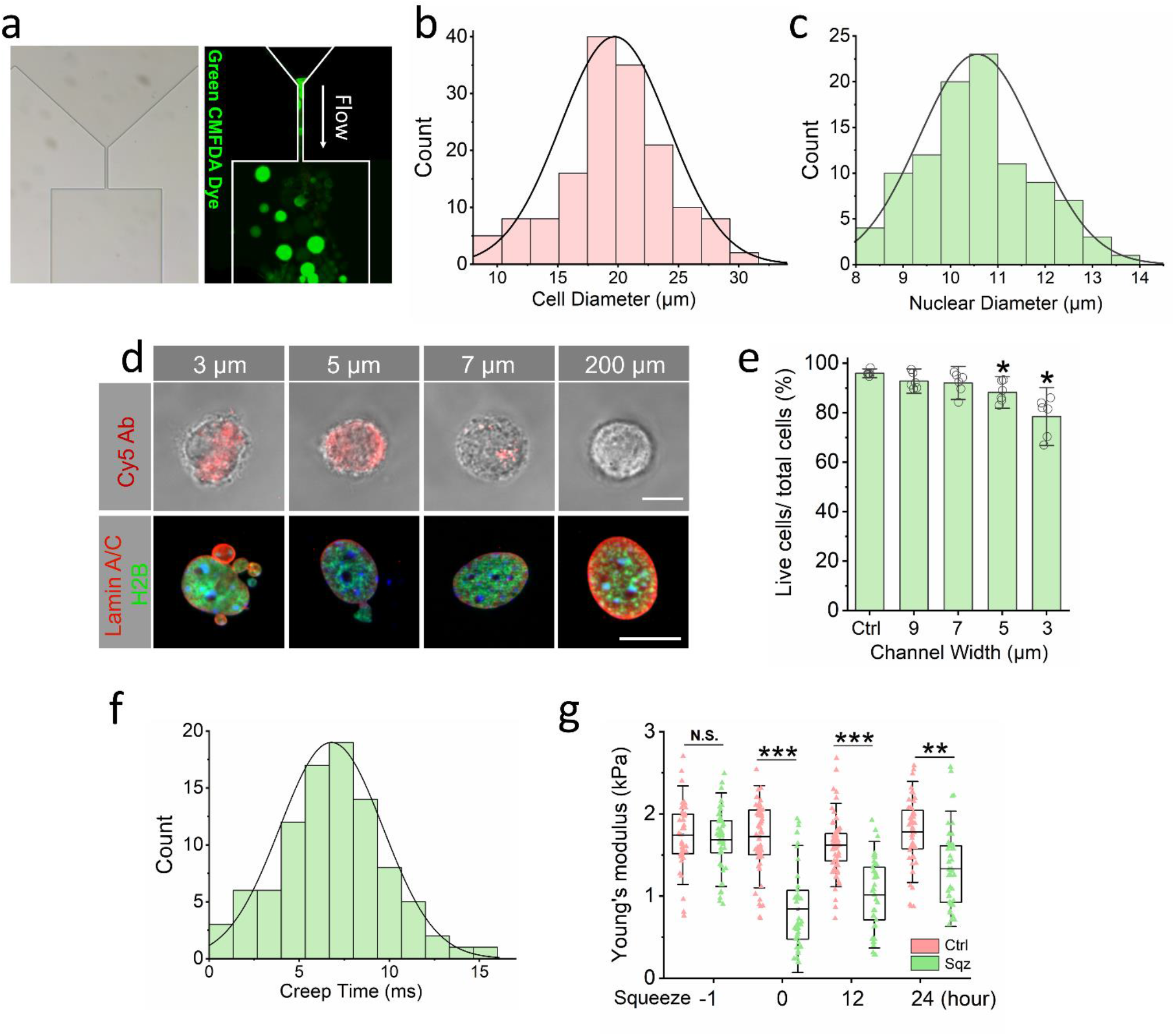
Microchannel-induced nuclear deformation. **(a)** Phase contrast and immunofluorescent images of microchannels (left) and fibroblasts labeled with Cell Tracker Green passing through the microchannels (right). **(b)** Quantification of cell diameter based on TC20 Cell Counter (Bio-Rad, USA) (n≥100). **(c)** Quantification of nuclear diameter based on Hoechst staining (Life technology, USA) (n≥100). **(d)** Phase contrast images of cells that were deformed by using microdevices with different widths of microchannels. Cell membrane rupture was detected immediately after cells passed through the microchannel using a Cy5-tagged antibody (top). Scale bar, 10 µm. Immunofluorescent images of H2B-GFP fibroblasts that were deformed by using microdevices with different widths of microchannel and seeded onto fibronectin-coated glass slides for 3 hours, and stained with lamin A/C. Scale bar, 10 µm. **(e)** The percentage of live cells after BAM-transduced fibroblasts were deformed using microdevices with various channel widths. Bar graphs show mean ± standard deviation (n=6). *p≤0.05 compared to the control. Statistical significance was determined by a one-way ANOVA and Tukey’s multiple comparison test. **(f)** Quantification of creep time in milliseconds as cells pass through the microchannels (n≥100). **(g)** Quantification of Young’s modulus of cells at the indicated time points before and after mechanical squeezing, as measured by AFM. Box plots show mean ± standard deviation (n≥50; **p≤0.01, ***p≤0.001), and significance was determined by a two-tailed, unpaired t-test at each time point.

As flow rate could affect the rate of forced nuclear deformation and cell aggregation in microchannels, we examined cell viability and channel clogging at various flow rates in 7-µm microchannels. As shown in **Supplementary Fig. S5**, the cell viability was not affected at a flow rate of 10 or 20 µL/minute. However, flow rates higher than 40 µL/min significantly decreased cell viability possibly due to the aggregation of cells in a single channel that caused clogging. Therefore, we used 20 µL/minute as an optimal flow rate to induce nuclear deformation. It took an average of 6.8 milliseconds (ms) for a cell introduced into a 7 µm-wide microchannel at 20 µL/minute flow rate, to pass through the microchannel (**Fig. 1f)**, which resulted in transient nuclear deformation.

To evaluate the effects of nuclear deformation on the mechanical properties of the nucleus, atomic force microscopy (AFM) was performed to measure the elastic modulus of cells at multiple time points after deformation. As shown in **Fig. 1g**, compared to control cells, the stiffness of the cells, measured by indenting the plasma membrane of the cells above the nucleus, decreased significantly after cells were squeezed through the channel. Over 24 hours, we observed a slight recovery indicated by increasing elastic modulus, indicating that the mechanical changes induced by transient squeezing lasted for hours. Additionally, we investigated nuclear shape following deformation by performing fluorescence imaging of cells stained with DAPI. We found that the cell nucleus was more elongated after squeezing, even after 24 hours (**Supplementary Fig. S6**).

To determine whether microchannel-induced nuclear deformation had any effect on the direct conversion of fibroblasts into iN cells, adult mouse fibroblasts were transduced with doxycycline (Dox)-inducible lentiviral constructs containing the three reprogramming factors BAM as depicted in the timeline for the reprogramming experimental procedure (**Fig. 2a**). The following day, Dox was added (designated as Day 0) to induce BAM expression, and BAM proteins were expressed within a few hours. Before or after administering Dox, cells were introduced into microfluidic devices with 7 μm-wide channels. Then the cells were collected and seeded onto fibronectin-coated glass coverslips and cultured in serum-free N2B27 medium (**Fig. 2a**). Seven days after induced mechanical deformation, cultures were fixed and stained for Tuj1 to determine the reprogramming efficiency. Administering Dox at least 6 hours before subjecting the cells to nuclear deformation produced the highest reprogramming efficiency (∼8-fold) compared to the control (**Fig. 2b**), suggesting that the presence of BAM proteins within the first few hours after squeezing was critical. Taken together, these results indicate that a transient nuclear deformation was sufficient to enhance the reprogramming efficiency of fibroblasts into iN cells. In addition, an increase in microchannel width to 9 µm did not further improve reprogramming efficiency (**Supplementary Fig. S7**).

**Fig. 2.**
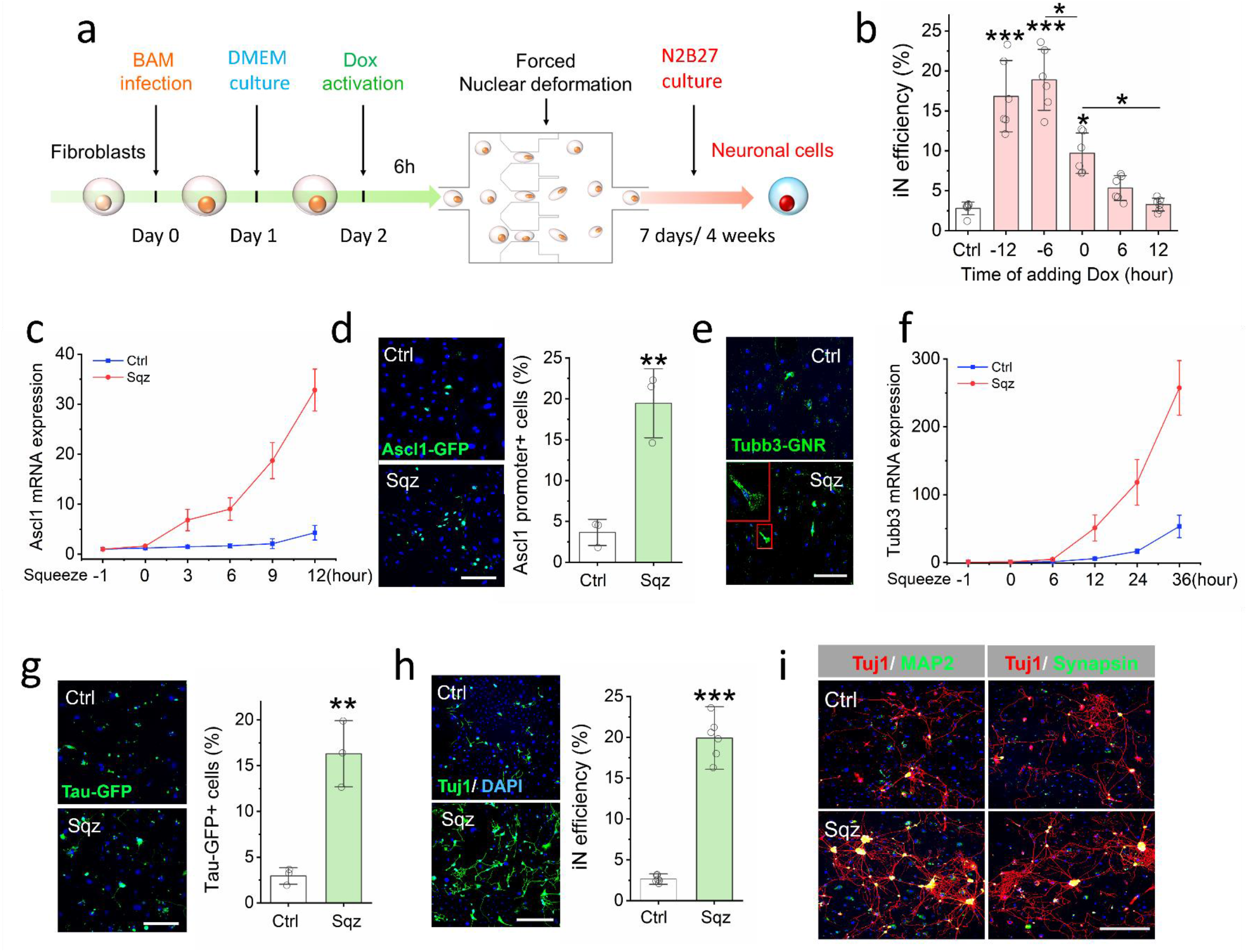
Microchannel-induced nuclear deformation accelerated the activation of neuronal marker expression and enhanced iN reprogramming efficiency. **(a)** Experimental timeline for nuclear deformation-induced iN reprogramming. **(b)** Timing effect of Dox administration on microchannel-induced iN reprogramming. Dox was added to BAM-transduced fibroblasts at the indicated time points before or after mechanical squeezing, and the reprogramming efficiency was determined at day 7. Bar graph shows mean ± SD (n=6, *p<0.05, ***p<0.001). Statistical significance was determined by a one-way ANOVA and Tukey’s multiple comparison test. **(c)** Relative Ascl1 mRNA expression in mechanically deformed and control cells at the indicated time points after nuclear deformation. Line graphs show mean ± standard deviation (n=6). **(d)** Fibroblasts transduced with BAM and an Ascl1 promoter-GFP construct were either mechanically deformed (squeezed: sqz) or kept as a control. The cells were fixed after 1 day, and observed by immunofluorescence microscopy, showing that squeezing induced more Ascl1 promoter-GFP^+^ cells at day 1, as quantified in the bar graph. Scale bar, 200 µm. Bar graphs show mean ± standard deviation (n=3; **p≤0.01), and significance was determined by a two-tailed, unpaired *t* test. **(e)** Immunofluorescent images show Tubb3 mRNA in mechanically deformed and control cells at 12 hours after mechanical squeezing, as detected by using a Tubb3 mRNA golden nano-rod (GNR) biosensor. Scale bar, 100 µm. **(f)** Relative Tubb3 mRNA expression in mechanically deformed and control cells at the indicated time points after nuclear deformation. Line graphs show mean ± standard deviation (n=6). **(g)** BAM-transduced Tau-GFP fibroblasts were either mechanically deformed or kept as a control, and cultured for 7 days. Immunofluorescent images show Tau-GFP^+^ cells at day 7, as quantified in the bar graph. Scale bar, 200 µm. Bar graphs show mean ± standard deviation (n=3; **p≤0.01) and significance was determined by a two-tailed, unpaired *t* test. **(h)** Reprogramming efficiency of BAM-transduced fibroblasts that were either deformed or kept as a control. At day 7, the cells were fixed and stained for Tuj1, followed by immunofluorescence microscopy to quantify Tuj1^+^ iN cells. Scale bar, 200 µm. Bar graphs show mean ± standard deviation (n=6; ***p≤0.001) and significance was determined by a two-tailed, unpaired *t* test. **(i)** Representative images of Tuj1^+^ cells expressing mature neuronal markers, MAP2 and synapsin at 4 weeks after nuclear deformation. Scale bar, 200 µm.

To determine whether nuclear deformation by microchannels accelerated iN conversion, we monitored the effect of nuclear deformation on neuronal marker activation by quantitative polymerase chain reaction (qPCR) and live cell imaging. While Ascl1 expression showed a basal level in the control group (likely transgene expression) and a slight increase at 12 hours (potential activation of endogenous Ascl1), squeezing cells triggered an early increase of Ascl1 expression at 3 hours, which continued to rapidly increase, resulting in a 7.6-fold increase when compared to the control group (**Fig. 2c**). In addition, to directly monitor the temporal activation of endogenous Ascl1, fibroblasts were transduced with an Ascl1 promoter driven-GFP (Ascl1-GFP) and subjected to reprogramming. Consistently, we found a significant increase (∼5-fold) in the number of Ascl1-GFP^+^ cells at day 1 in the squeezed cells compared to the control (**Fig. 2d**). These results suggest that nuclear deformation not only activated neuronal gene expression earlier, but also enhanced reprogramming efficiency by activating neuronal genes in more cells.

Furthermore, we monitored the expression of other neuronal markers such as neuronal beta-III tubulin (*Tubb3*). We first used gold nanorod biosensors^26^ with complementary sequence to detect mRNA expression of *Tubb3* in living cells. *Tubb3* mRNA expression was detectable as early as 12 hours in squeezed cells but rarely in the control group (**Fig. 2e, Supplementary Fig. S8**). Consistently, quantitative reverse transcription-polymerase chain reaction (qRT-PCR) analysis showed significantly higher *Tubb3* expression in the squeezed cells than control cells at multiple time points after nuclear deformation (**Fig. 2f**).

In addition, we performed reprogramming experiments using fibroblasts isolated from transgenic mice expressing a EGFP reporter driven by the promoter of neuronal gene Tau. The number of Tau-GFP^+^ cells in the squeezed group at day 4 was ∼6 times greater than that in the control group in these cells as well (**Fig. 2g**). At 1 week and 2 weeks after nuclear deformation, the iN reprogramming efficiency was significantly higher in the mechanically squeezed group compared to the control (**Fig. 2h, Supplementary Fig. S9**). Further characterization of the derived cells revealed that iN cells expressed mature neuronal markers microtubule associated protein 2 (MAP2) and synapsin at 4 weeks after nuclear deformation (**Fig. 2i**). After 6 weeks, iN cells resulted from mechanical squeezing treatment showed calcium fluctuation, indicating a mature neuronal phenotype (**Supplementary movie S1**)^27^, suggesting that although mechanical squeezing promoted reprogramming, it did not interfere with the neural differentiation process.

The cell reprogramming process involves a significant reorganization of chromatin and changes in the epigenetic state that controls the on/off state of phenotypic genes. To investigate whether nuclear deformation-enhanced reprogramming efficiency was due to changes in chromatin reorganization and the epigenetic state, we first utilized a fluorescence resonance energy transfer (FRET) biosensor targeted at the nucleosome to monitor the change in levels of a heterochromatin mark H3K9me3. To observe the change in single cells, we slowed down the squeezing process by lowering the pressure and flow rate in the microdevice. We found that H3K9me3 FRET signal significantly decreased in fibroblasts passing through the microchannels (**Supplementary Fig. S10**), indicating that the transient nuclear deformation induced an essentially concurrent change in this histone mark. To determine whether the epigenetic changes persisted after squeezing and whether squeezing induced changes in other epigenetic marks, we performed immunostaining analysis of euchromatin and heterochromatin marks at multiple time points within the first 24 hours after cells passed through the microchannels. Consistently, we observed a significant decrease in H3K9me3 at 3 hours and 12 hours after nuclear deformation, which return to the same level as the cells in the control group after 24 hours (**Fig. 3a-b and Supplementary Fig. S11**), suggesting that nuclear deformation resulted in a transient reduction of heterochromatin. In contrast, the levels of acetylated histone marks including ACH3 and H3K9ac, and specific histone methylation marks including H3K4me1, H4K20me3 and H3K27me3 did not show significant global changes in response to forced nuclear deformation (**Supplementary Fig. S12**).

**Fig. 3.**
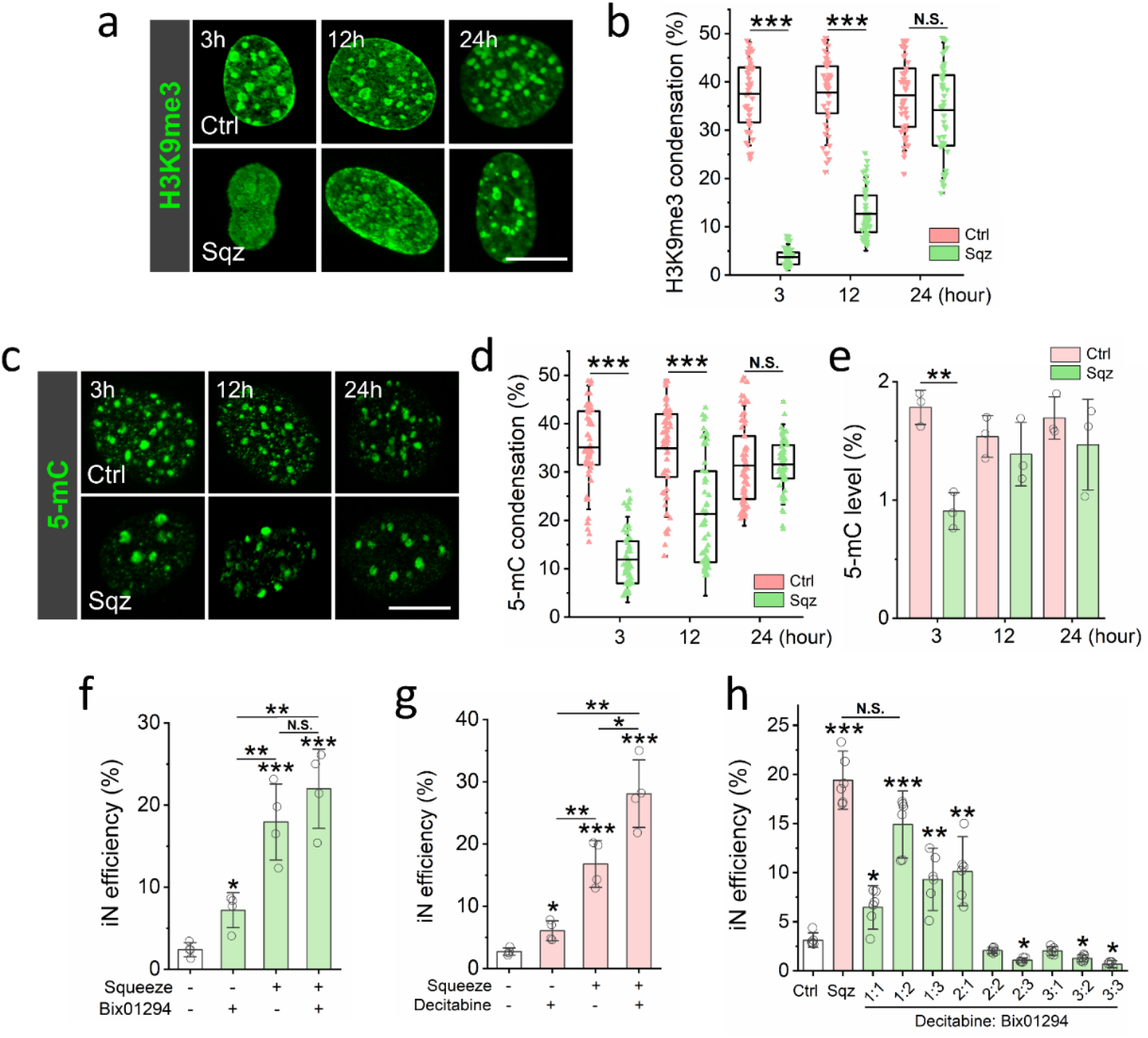
Nuclear deformation induced the demethylation of histone and DNA. **(a)** Immunostaining of H3K9me3 in the nucleus at the indicated time points after mechanical squeezing. Scale bar, 10 µm. **(b)** Quantification of H3K9me3 condensation in control and squeezed cells at the indicated time points (based on experiments in **a**). Box plots show mean ± standard deviation (n ≥ 40; ***p≤0.001), and significance was determined by a two-tailed, unpaired *t* test for each time point. **(c)** Representative images of 5-mC staining in control and squeezed cells at various time points. Scale bar, 10 µm. **(d)** Quantification of 5-mC condensation in control and squeezed cells at the indicated time points (based on experiments in **c**). Box plots show mean ± standard deviation (n ≥ 40; ***p≤0.001), and significance was determined by a two-tailed, unpaired *t* test for each time point. **(e)** Quantification of 5-mC level in control and squeezed cells at the indicated time points. Bar graphs show mean ± standard deviation (n=3; *p≤0.05), and significance was determined by a two-tailed, unpaired *t* test for each time point. **(f)** Reprogramming efficiency of BAM-transduced fibroblasts that were cultured in the absence or presence of 1 μM Bix01294 and stimulated by the microfluidic device. Bar graphs show mean ± standard deviation (n=4; *p≤0.05, **p≤0.01, ***p≤0.001), and significance was determined by a one-way ANOVA and Tukey’s multiple comparison test. **(g)** Reprogramming efficiency of BAM-transduced fibroblasts in the presence or absence of 0.5 μM Decitabine and stimulated by the microfluidic device. Bar graphs show mean ± standard deviation (n=4; *p≤0.05, **p≤0.01, ***p≤0.001), and significance was determined by a one-way ANOVA and Tukey’s multiple comparison test. **(h)** Reprogramming efficiency of BAM-transduced fibroblasts treated with different amounts of Bix01294 and Decitabine. 1: 0.2 μM; 2: 0.4 μM; 3: 0.6 μM. Bar graphs show mean ± standard deviation (n=6; *p≤0.05, **p≤0.01, ***p≤0.001), and significance was determined by a one-way ANOVA and Tukey’s multiple comparison test.

In addition to histone modifications, DNA methylation influences chromatin organization, which is critical for cell reprogramming^28^. To investigate the effect of nuclear deformation on DNA methylation, we analyzed DNA condensation and the level of 5-methylcytosine (5-mC), a DNA methylation marker, in fibroblasts squeezed by microchannels. As shown in **Fig. 3c-e** and **Supplementary Fig. S13**, nuclear deformation significantly decreased DNA methylation for at least 12 hours, suggesting that nuclear deformation may affect chromatin structure through changes in both DNA and histone methylation.

To test whether the decrease in H3K9me3 played a role in iN conversion, BAM-transduced fibroblasts were treated with a H3K9 specific methyltransferase inhibitor (Bix01294) for 24 hours. The inhibitor suppressed H3K9me3 in a dose-dependent manner, and we selected a concentration of Bix01294 (1 μM) that did not affect cell viability (**Supplementary Fig. S14-15**). As shown in **Fig. 3f**, Bix01294 partially mimicked the squeezing effect on the reprogramming efficiency. Additionally, pre-treatment with Bix01294, together with mechanical deformation, did not further increase the reprogramming efficiency compared to the squeezed-only groups (**Fig. 3f)**. Furthermore, to determine whether the decrease of H3K9me3 was required for nuclear deformation-induced iN reprogramming, BAM-transduced fibroblasts were pre-treated with JIB-04 (100 nM), a H3K9 specific histone demethylase inhibitor, for 24 hours before being introduced into the microdevice. JIB-04, which significantly increased H3K9me3, not only reduced the reprogramming efficiency compared to the control, but also strikingly suppressed nuclear deformation-induced iN reprogramming (**Supplementary Fig. S16**). These results suggest that the decrease in H3K9me3 levels was required for the mechanical squeezing effect on iN reprogramming. Pre-treatment with the DNA methyltransferase inhibitor, Decitabine (0.5 μM), slightly enhanced iN reprogramming efficiency; pre-treatment with Decitabine before squeezing further enhanced squeezing-induced iN reprogramming (**Fig. 3g, Supplementary Fig. S14-15**), demonstrating that DNA demethylation also played a role in iN reprogramming.

To investigate whether the combined effects of suppressing H3K9me3 and DNA methylation could match the reprogramming efficiency induced by mechanical squeezing, BAM-transduced fibroblasts were either subjected to transient nuclear deformation or treated with different combinations and concentrations of methyltransferase inhibitors for H3K9 and DNA. As shown in **Supplementary Fig. S17 and S18**, both Bix01294 and Decitabine significantly increased iN conversion. Interestingly, 0.2 μM Decitabine combined with 0.4 μM Bix01294 (1:2) significantly increased the iN reprogramming efficiency compared to the other combinations tested and showed similar efficiency when compared with the mechanically squeezed group (**Fig. 3h**), suggesting that the suppression of H3K9me3 and DNA methylation may be the major mediators of microchannel-induced iN reprogramming efficiency.

We further investigated whether ion channels were involved in squeezing-induced mechanotransduction leading to improved reprogramming efficiency. As shown in **Supplementary Fig. S19-23**, the inhibition of Na^+^, K^+^, and Ca^2+^ ion channels and the manipulation of extracellular pH did not significantly affect the iN reprogramming efficiency after transient nuclear deformation.

The observations that mechanical squeezing forced nuclear deformation and a significant decrease in Young’s modulus of the cells and altered nuclear shape suggested that the structural changes of the nucleus could mediate mechanotransduction through the nuclear matrix. Indeed, lamin A/C staining showed a transient decrease in lamin assembly at the nuclear periphery and a transient increase in the wrinkling of the nuclear membrane for at least 6 hours (**Fig. 4a-c and Supplementary Fig. S24**). In addition, the structure of the lamin A/C was disrupted within 1 hour after transient nuclear deformation compared to the control cells (**Supplementary Fig. S25**). Further examination of H3K9me3 and 5-mC revealed that H3K9me3 was no longer co-localized with lamin A/C at the nuclear periphery, and there was a disassembly of 5-mC clusters (**Supplementary Fig. S25**). To investigate whether lamin disruption mediated nuclear deformation-induced epigenetic changes during iN reprogramming, we silenced lamin A by using a small interfering RNA (siRNA) in BAM-transduced fibroblasts 24 hours prior to squeezing with microchannels (**Supplementary Fig. S26**). As shown in **Fig. 4d**, lamin A knockdown mimicked the effects of mechanical squeezing, which caused a lamin disassembly and a decrease in H3K9me3 and 5-mC (**Fig. 4e, f**). Moreover, lamin knockdown enhanced the iN reprogramming efficiency of non-deformed cells to an extent similar to the cells that were mechanically deformed but lamin A was not silenced (**Fig. 4g**). The iN cells derived after lamin A knockdown expressed MAP2 and synapsin at 4 weeks after mechanical deformation (**Supplementary Fig. S27**). Taken together, these results suggested that lamin A played an important role in regulating the demethylation of histone and DNA induced by forced nuclear deformation.

**Fig. 4.**
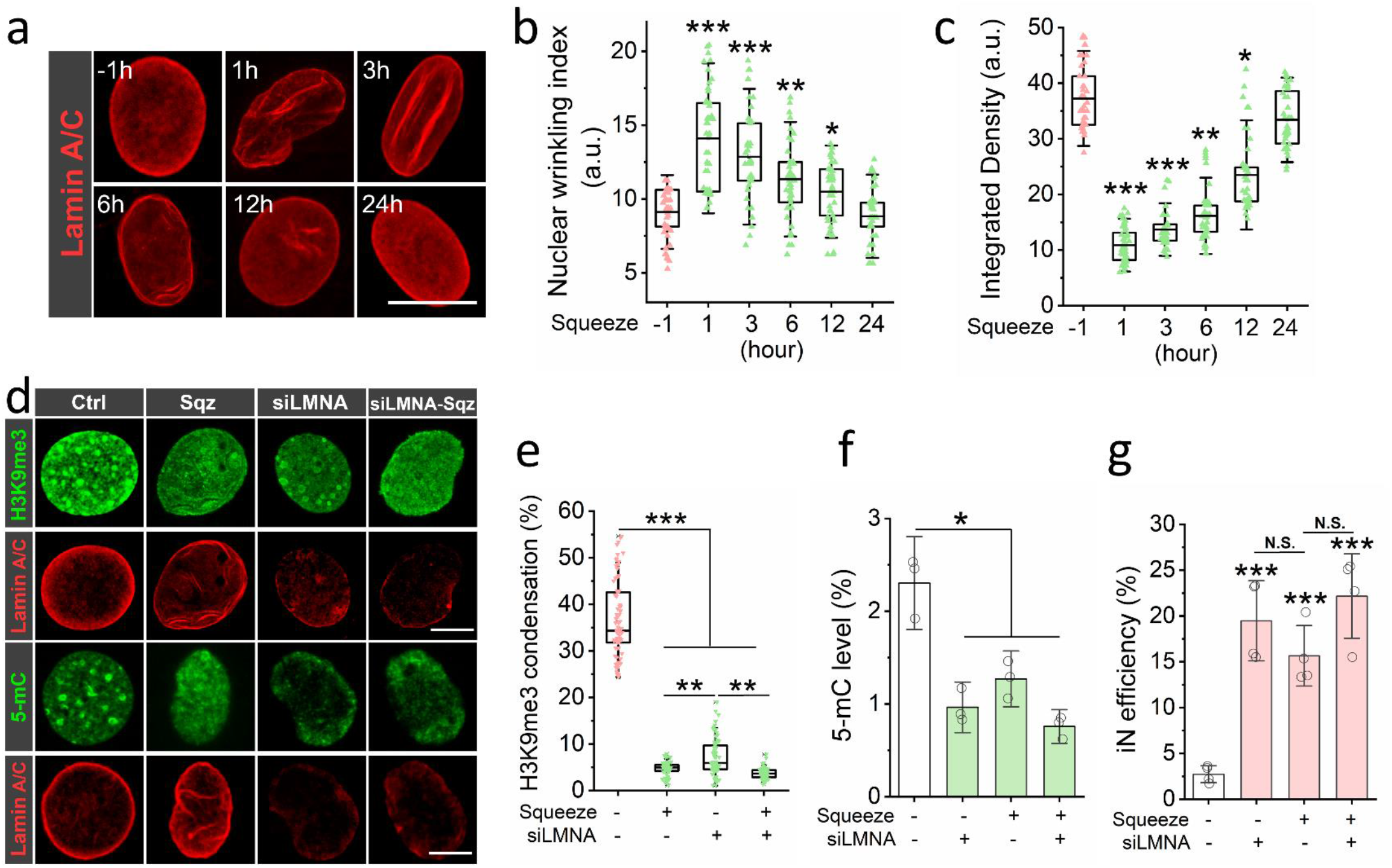
Lamin A/C mediated microchannel-induced epigenetic changes and iN reprogramming. **(a)** Representative images of lamin A/C staining in squeezed cells at the indicated time points after mechanical deformation. Scale bar, 10 µm. **(b)** Quantification of nuclear wrinkling index at the indicated time points before and after mechanical squeezing. Nuclear wrinkling was defined as the immunofluorescence intensity variation within the nuclear region. Bar graphs show mean ± standard deviation (n≥50; *p≤0.05, **p≤0.01, ***p≤0.001) and significance was determined by a one-way ANOVA and Tukey’s multiple comparison test. **(c)** Quantification of the integrated density of lamin A/C at the indicated time points before and after mechanical squeezing. Bar graphs show mean ± standard deviation (n≥50; *p≤0.05, **p≤0.01, ***p≤0.001), and significance was determined by a one-way ANOVA and Tukey’s multiple comparison test. **(d)** Immunostaining of H3K9me3 and 5-mC in control and lamin A-silenced fibroblasts at 3 hours after mechanical squeezing. Scale bar, 5 µm. **(e)** Quantification of H3K9me3 condensation in control and squeezed cells after lamin A knockdown (based on experiments in **d**). Box plots show mean ± standard deviation (n ≥ 50; *p≤0.05, **p≤0.01, ***p≤0.001), and significance was determined by a one-way ANOVA and Tukey’s multiple comparison test. **(f)** Quantification of 5-mC in control and lamin A-silenced fibroblasts at 3 hours after mechanical squeezing. Bar graphs show mean ± standard deviation (n=3; *p≤0.05), and significance was determined by a one-way ANOVA and Tukey’s multiple comparison test. **(g)** Reprogramming efficiency of control and lamin A-silenced BAM-transduced fibroblasts stimulated by the microfluidic device. Bar graphs show mean ± standard deviation (n=4; ***p≤0.001) and significance was determined by a one-way ANOVA and Tukey’s multiple comparison test.

To determine whether microfluidic device-induced nuclear deformation could regulate the reprogramming of different cell types, we performed similar experiments using macrophages transduced with BAM and fibroblasts transduced with Oct-4, Sox 2, KLF-4, and c-Myc (OSKM), respectively. Interestingly, we found that both iN reprogramming from macrophages and iPSC reprogramming from fibroblasts were significantly enhanced after nuclear deformation (**Fig. 5a, b**), suggesting a general approach to mechanically modulate the epigenetic state of fibroblasts or other cell types to promote cell reprogramming.

**Figure 5.**
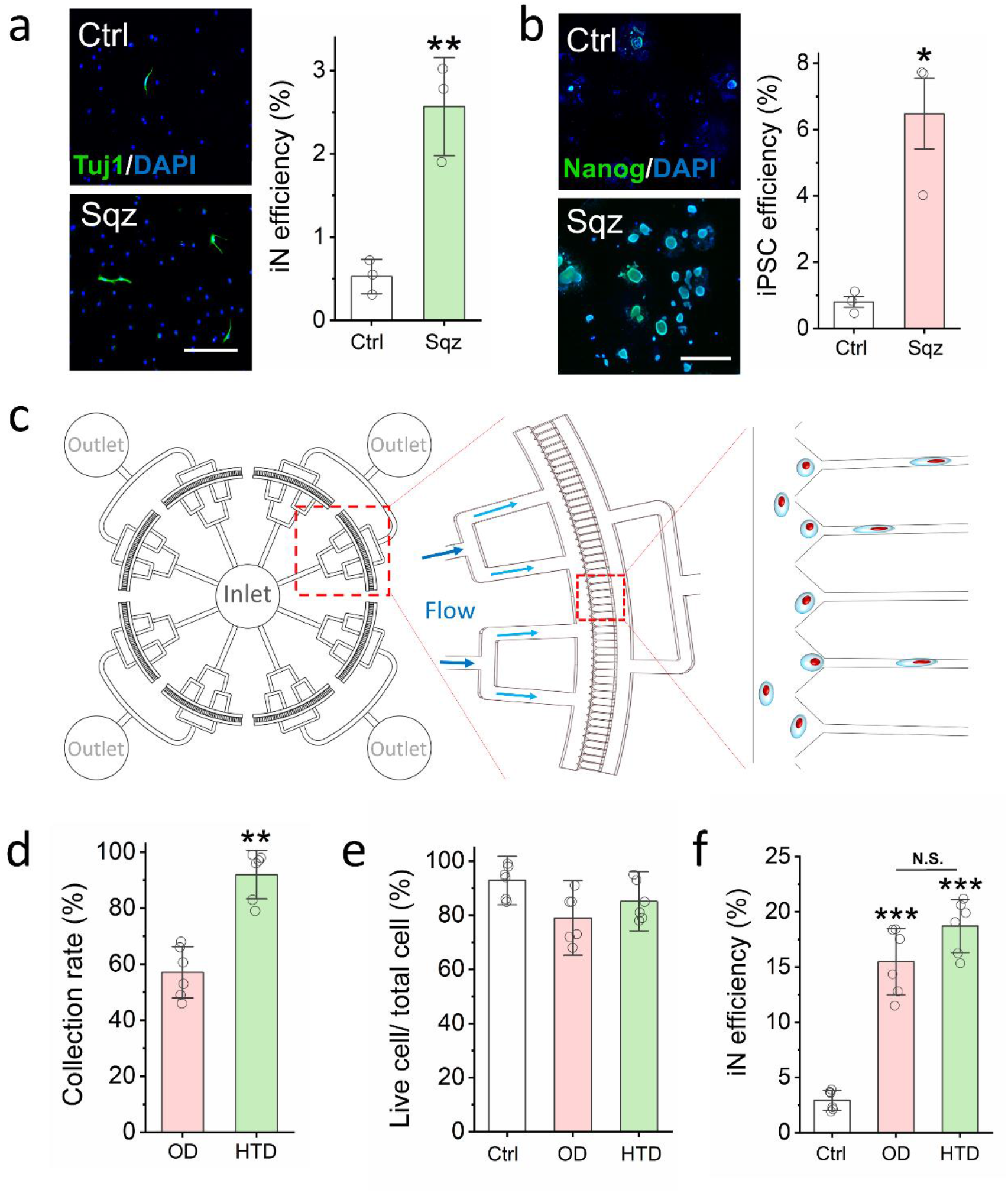
Development of cell reprogramming technologies by nuclear deformation. **(a)** Immunofluorescent images show Tuj1^+^ iN cells generated from BAM-transduced mouse macrophages that were stimulated with the microfluidic device. Scale bar, 200 μm. Reprogramming efficiency of BAM-transduced macrophages after mechanical squeezing. Bar graph shows mean ± SD (n=3, **p<0.01). Statistical significance was determined by a two-tailed, unpaired t-test. **(b)** Immunofluorescent images show Nanog^+^ colonies generated from OSKM-transduced mouse fibroblasts that were stimulated with the microdevice. Scale bar, 1 mm. Reprogramming efficiency of OSKM-transduced mouse fibroblasts into iPSCs after mechanical squeezing. Bar graph shows mean ± SD (n=3, *p<0.05). Statistical significance was determined by a two-tailed, unpaired t-test. **(c)** Schematic illustrating the design of the high-throughput device. **(d)** Quantification of the percentage of cells that were collected after passing through the original microchannel device (OD) or high-throughput microfluidic device (HTD). Bar graph shows mean ± SD (n=6, **p<0.01). Statistical significance was determined by a two-tailed, unpaired t-test. **(e)** The percentage of live cells after BAM-transduced fibroblasts were deformed using OD or HTD device. Bar graphs show mean ± standard deviation (n=6) and significance was determined by a one-way ANOVA and Tukey’s multiple comparison test. **(f)** Reprogramming efficiency of BAM-transduced fibroblasts that were either kept as a control or deformed by using the OD or HTD microfluidic device at day 7. Bar graphs show mean ± standard deviation (n=6; ***p≤0.001) and significance was determined by a one-way ANOVA and Tukey’s multiple comparison test.

To scale up the mechano-preconditioning of cells for reprogramming, we developed a higher-throughput device containing 10 times more microchannels (400 microchannels) than the original device with 36 microchannels (**Fig. 5c)**. The design was validated by fluid flow simulation, showing the velocity profile in different sizes of channels in the devices **(Supplementary Fig. S28**). By using the higher-throughput device, 10,000 fibroblasts could be collected within a minute, which was 5 times faster than the original device. Compared to our original device (OD) (**Supplemental Figure 1a**), this high-throughput device (HTD) significantly increased the yield of cell collection, with more than 95% of cells introduced into device collected at the outlets (**Fig. 5d**). Additionally, this high-throughput device maintained cell viability and significantly improved iN efficiency (**Fig. 5e, f**).

## Discussion

In this study, we demonstrate that the transient nuclear deformation in suspended cells decreases the methylation of H3K9me3 and DNA (5-mC), which primes the chromatin to a more permissive epigenetic state for reprogramming. This phenomenon is independent of complex microenvironmental factors such as extracellular matrix, cell-cell adhesions and pH, and may be generalized to various cell types as we have shown for fibroblasts and macrophages. It is important to note that this millisecond mechanical perturbation induces a transient change in epigenetic state within a 24-hour time window, allowing an earlier and more efficient activation of neuronal genes in heterochromatin, with global heterochromatin marks returning to the basal level afterwards. It is also interesting that this simple mechanical squeezing is more effective than chemicals that cause the demethylation of H3K9 and DNA (5-mc), suggesting that nuclear deformation may induce many changes in chromatin and/other signaling events that facilitate cell reprogramming.

The simplicity of this approach by squeezing suspended cells provides a direct evidence on the effect of nuclear deformation on chromatin remodeling. Passive biophysical factors such as micro/nano topography and matrix stiffness not only cause nuclear deformation, but also induce changes in cell adhesions and cytoskeleton organization that affect many other cellular processes. Active mechanical loading such as magnetic twisting on the surface of adherent cells, although regulating gene expression in euchromatin, appears insufficient to overcome the barrier heterochromatin^29^, which could be explained by the lack of significant nuclear deformation and/or chromatin reorganization. Stretching or compressing adherent cells may decrease or increase heterochromatin^21–23^, and these different effects could be related to the different magnitudes and rates of nuclear perturbation by these mechanical stimulations and other confounding factors such as cell adhesions and polarity.

FRET experiments show that nuclear deformation downregulates H3K9me3 within minutes, which is accompanied by lamin disassembly and a decrease in cell stiffness. This is consistent with an earlier observation during cell mitosis^30^. Knocking down lamin A mimics the effects of mechanical squeezing, suggesting that the nuclear matrix plays an important role in this mechanotransduction process. Indeed, heterochromatin is anchored to the lamin A-associated domains that are abundant for heterochromatin marks such as H3K9me3 and are repressive for gene expression^31–33^. Mechanical force-induced disruption of nucleus may loosen heterochromatin positioned at the nuclear periphery and relocate it towards the interior of the nucleus, accompanied by the downregulation of heterochromatin marks. This transient biophysical modulation of epigenetic state appears to be universal and independent of cell type and reprogramming factors, as squeezing macrophages also enhances their reprogramming into neurons and the transient nuclear deformation promotes fibroblast reprogramming into iPSCs.

Another highlight of this work is the translation of mechanobiology findings into mechano-biotechnology for cell engineering. Microfabricated devices provide a well-controlled microenvironment and real-time process control with a minimal benchtop space requirement^34^. Here we developed a scalable microfluidic device that can be used to continuously process and precondition cells (approximately 10,000 cells per minute). This microfluidic device with multiple constriction channels can be used to engineer a variety of cells such as fibroblasts, stem cells and immune cells, and facilitate the conversion of cell types from one to another, which will have broad applications in regenerative medicine, disease modeling, and drug screening.

## Methods

### Microfabrication of the microfluidic device

The molds of designed microfluidic devices for cell squeezing were fabricated via photolithography. A 15-µm thick layer of SU-8 2015 (Microchem Corporation, 3300 rpm) was spun coated onto a 4-inch silicon wafer, followed by standard photolithography process according to the manufacturer’s instruction. Base and curing agent of polydimethylsiloxane (PDMS, Sylgard 184, Dow Corning) was mixed at a 10:1 weight ratio and degassed in a vacuum chamber for one hour to remove air bubbles before being poured onto the mold. After curing at 65 °C for 4 hours, the PDMS mold was punched to make inlets and outlets for tubing connections. The PDMS mold and pre-cleaned glass were bonded after treatment with oxygen plasma for 30 seconds. The bonded chips were baked at 65 °C for 10 minutes to enhance the bonding.

### Cell isolation, culture and reprogramming

Fibroblasts were isolated from ear tissues of adult (1 month-old) C57BL/6 mice, Tau-EGFP reporter mice (Jackson Laboratory, 004779) and R26-M2rtTA;Col1a1-tetO-H2B-GFP compound mutant mice (Jackson Laboratory, 016836), and expanded in fibroblast medium: DMEM (Gibco, 11965), 10% fetal bovine serum (FBS; Gibco, 26140079) and 1% penicillin/streptomycin (GIBCO, 15140122). For all experiments, passage-2 cells were used and synchronized upon reaching 80% confluency using DMEM with 1% FBS for 24 hours before the transduction with viruses containing BAM constructs. The following day (day 0) the medium was changed to mouse embryonic fibroblasts (MEF) medium containing doxycycline (2 ng/ml, Sigma) to initiate the expression of the transgenes and thus, reprogramming. After 6 hours, transduced fibroblasts were passaged, and either subjected to microfluidic deformation or kept as controls. Cells were then seeded onto glass slides coated with 0.1 mg/mL fibronectin (ThermoFisher, 33016015) overnight at a density of 3,000 cells/cm^2^. Twenty-four hours later (day 1), cells were cultured in N2B27 medium: DMEM/F12 (Gibco, 11320033), N-2 supplement (Gibco, 17502048), B-27 supplement (Gibco, 17504044), 1% penicillin/streptomycin, and doxycycline (2ng/ml), and half medium changes were performed every 2 days. On day 7 after microfluidic deformation, cells were fixed and stained for Tuj1 to determine the reprogramming efficiency. iN cells were identified based on positive Tuj1 staining and a neuronal morphology. The reprogramming efficiency was determined as the percentage of iN cells on day 7 relative to the number of the cells initially seeded. For long-term studies where maturation and functionality of the iN cells were examined, cells were kept in culture for 5 weeks. Reprogramming of iPSC from wild-type fibroblasts was performed as described previously^18^.

Macrophages for reprogramming experiments were derived from differentiated monocytes. Monocytes were isolated from the bone marrow of adult C57BL/6 mice, and expanded in monocyte medium: RPMI 1640 (Gibco, 11875093), 10% fetal bovine serum (FBS; Gibco, 26140079) and 1% penicillin/streptomycin (GIBCO, 15140122). The next day, macrophage-colony stimulating factor (M-CSF) (50 ng/ml, ThermoFisher, PMC2044) was added to the medium and cells were cultured for an additional 2 days. Cells were then washed 3 times with phosphate buffered saline (PBS) before transduction with viruses containing BAM constructs.

### Lentiviral preparation and transduction

Doxycycline-inducible lentiviral vectors for Tet-O-FUW-Brn2, Tet-O-FUW-Ascl1, Tet-O-FUW-Myt1l, Tet-O-FUW-EGFP, and FUW-rtTA plasmids were used to transduce fibroblasts for ectopic expression of Brn2, Ascl1, Myt1L, GFP, and rtTA. The STEMCCA lentiviral vector was used for the ectopic expression of OSKM^18^. The Ascl1-eGFP lentiviral vector (Genecopoeia, MPRM39894-LvPF02) was used to monitor the activation of the Ascl1 promoter. Lentivirus was produced by using established calcium phosphate transfection methods, and Lenti-X Concentrator (Clontech, 631232) was utilized to concentrate viral particles according to the manufacturer’s protocol. Stable virus was aliquoted and stored at − 80°C. Fibroblasts were plated and synchronized for 24 hours before viral transduction in the presence of polybrene (8µg/ml; Sigma, H9268). Cells were incubated with the virus for 24 hours before performing microfluidic deformation experiments.

### Cell viability assays

After cells passed through the micro-device, 10×10^3^ fibroblasts were plated and allowed to attach for 3 hours in 96 well plate. Live and dead assays were performed using the LIVE/DEAD™ Cell Imaging Kit (Invitrogen, R37601) according to the manufacturer’s protocol. Cells were incubated with an equal volume of 2X working solution for 15 minutes at room temperature. Epifluorescence images were collected using a Zeiss Axio Observer Z1 inverted fluorescence microscope and analyzed using ImageJ.

Cell viability was assayed using the PrestoBlue® Cell Viability Reagent (Invitrogen, A13261) according to the manufacturer’s protocol. Cells were incubated with the PrestoBlue Reagent for 2 hours. Absorbance was measured by a plate reader (Infinite 200PRO) at excitation/emission= 560/590 nm. Results were normalized to control (i.e., cell passing through >200 µm channels) samples.

### Immunofluorescence staining and microscopy

Samples collected for immunofluorescence staining at the indicated time points were washed once with PBS and fixed in 4% paraformaldehyde for 15 minutes. Samples were washed three times with PBS for 5 minutes each and permeabilized using 0.5% Triton X-100 for 10 minutes. After three subsequent PBS washes, samples were blocked with 5% normal donkey serum (NDS; Jackson Immunoresearch, 017000121) in PBS for 1 hour. Samples were incubated with primary antibodies (**Supplementary Table S1**) in antibody dilution buffer (1% normal donkey serum (NDS) + 0.1% Triton X-100 in PBS) for either 1 hour or overnight at 4°C followed by three PBS washes and a 1-hour incubation with Alexa Fluor® 488- and/or Alexa Fluor® 546-conjugated secondary antibodies (Molecular Probes). Nuclei were stained with DAPI in PBS for 10 minutes. Epifluorescence images were collected using a Zeiss Axio Observer Z1 inverted fluorescence microscope and analyzed using ImageJ. Confocal images were collected using a Leica SP8-STED/FLIM/FCS Confocal and analyzed using ImageJ.

For DNA methylation staining, samples were fixed with ice-cold 70% ethanol for 5 minutes followed by three PBS washes. Samples were then treated with 1.5M HCl for 30 minutes and washed thrice with PBS. The immunostaining procedure proceeded from the donkey serum blocking step as aforementioned.

### Chemical treatment of cells

To determine the role of H3K9 methylation on microfluidic device-induced iN reprogramming, BAM-transduced fibroblasts were treated with the H3K9 methyltransferase inhibitor Bix-01294 (Cayman chemical, 13124) or demethylase inhibitor JIB-04 (Cayman chemical, 15338) at the indicated concentrations for 24 hours prior to introduction into the microdevice. Parallel conditions with DMSO served as a control. The iN reprogramming efficiency was detected via Tuj1 staining 7 days after squeezing.

To determine the involvement of ion channels in microfluidic device-induced iN reprogramming, calcium channel blocker Amlodipine (Cayman chemical, 14838), potassium channel blocker Quinine (Cayman chemical, 23958) and sodium channel blocker procainamide (Cayman chemical, 24359) were used to inhibit calcium, potassium, and sodium ion channels, respectively. BAM-transduced fibroblasts were treated with small molecule blockers at the indicated concentrations for 12 hours prior to being introduced into the microdevice. Parallel conditions with DMSO served as a control. The iN reprogramming efficiency was detected via Tuj1 staining 7 days after squeezing.

To determine the effect of pH on forced nuclear deformation-induced iN reprogramming, BAM-transduced fibroblasts were treated with DMEM medium at different pH levels (pH=6.5, 7.5 and 8.5) for 1 hour prior to being introduced into the microdevice. The iN reprogramming efficiency was detected via Tuj1 staining at day 7 day after squeezing.

### DNA methylation assay

After cells passed through the device, cells were collected and 10×10^5^ cells were plated in 60 mm dishes. At different time points, cells were trypsinized and DNA was extracted by Invitrogen PureLink Genomic DNA mini kit (Invitrogen, K1820-01). The 5-mC level was analyzed by the MethylFlash™ Global DNA Methylation (5-mC) ELISA Easy Kit (Epigentek, P-1030) according to the manufacturer’s instructions. Briefly, 100 ng of sample DNA was bonded into the assay wells and incubated with a 5-mC detection complex solution for 60 minutes. Then color developer solution was added into assay wells, and the absorbance at 450 nm was measured by using a plate reader (Infinite 200Pro, 30050303).

### Quantitative reverse transcription polymerase chain reaction (qRT-PCR)

After cells passed through the device, cells were collected and 10×10^5^ cells were plated in 60-mm dishes. At different time points, TRIzol™ Reagent (Invitrogen, 15596026) was used to lyse cells, and RNA was isolated as described previously^35^. After RNA extraction, ThermoScientific Maxima First Strand cDNA Synthesis Kit (ThermoFisher, K1641) was used for first-strand cDNA synthesis. Then qRT-PCR was performed to detect the gene expression levels of *Ascl1* (Forward primer: GAAGCAGGATGGCAGCAGAT, Reverse primer: TTTTCTGCCTCCCCATTTGA) and *Tubb3* (Forward primer: GCGCCTTTGGACACCTATTC, Reverse primer: CACCACTCTGACCAAAGATAAAGTTGT), where 18S (Forward primer: GCCGCTAGAGGTGAAATTCTTG, Reverse primer: CATTCTTGGCAAATGCTTTCG) level was used for normalization.

### Lamin A siRNA knockdown

For Lamin A siRNA knockdown, 1 × 10^6^ cells were plated in 60-mm dishes for 24 hours. RNA interference was performed using ON-TARGETplus LMNA siRNA (Dharmacon, L-040758-00-0005), and transfections were carried out using Lipofectamine™ 3000 Reagent (ThermoFisher, L3000015) according to the manufacturer’s protocol. Briefly, 250 μl Opti-MEM™ Medium (ThermoFisher, 31985062) was mixed with 7.5 μl Lipofectamine™ 3000 Reagent and incubated at 37 °C for 15 minutes. At the same time, 5 μg siRNA was diluted in 250 μl Opti-MEM™ Medium and incubated at 37 °C for 15 minutes. These two solutions were mixed and the DNA-lipid complexes to cells were added to 1.5 ml DMEM medium without FBS and penicillin/streptomycin and incubated at 37 °C for 12 hours. The media was then replaced with DMEM medium with 10% FBS and 1% penicillin/streptomycin. Two days after transfection, RNA was isolated and qRT-PCR was performed to detect *LMNA* (Forward primer: GTCTCGAATCCGCATTGACA, Reverse primer: TGGCTGCCAACTGCTTTTG) mRNA expression to determine whether Lamin A had been silenced.

### Golden nanorod (GNR) LNA probe for mRNA

To detect Tubb3 mRNA expression in living cells after cells were passed through the device, cells were collected and plated in a 24 well plate at 2000 cells/well, and GNR-LNA complexes specific to Tubb3 was added to culture media as described previously^26^. Briefly, GNR-LNA complexes were made by mixing 1.5 μl LNA Probe (10 μM), 2.5 μl golden nano-rod (GNR) and 46 μl Tris-EDTA buffer, and incubating at 37°C for 15 minutes. The GNR-LNA complex solution (50 μl) and fresh culture medium (450 μl) were then mixed and added to the cells. After 4-hour incubation, cells were washed with PBS, and fresh culture medium was added. Cells were incubated at 37 ℃ in the dark for additional 60 minutes prior to performing live cell imaging. Epifluorescence images were collected using a Zeiss Axio Observer Z1 inverted fluorescence microscope.

### AFM measurement of cell mechanical property

To determine the elastic modulus of cells after passing through the device, mechanical measurements of single cells were performed by using atomic force microscopy (AFM) (JPK Nanowizard 4a) with tipless cantilevers (NPO-10, Bruker Corp., USA), a high sensitive cantilever k=0.06 N/m, and sample Poisson’s ratio of 0.499 at the UCLA Nano and Pico Characterization facility. During the measurement, cells were cultured on a glass-bottom dish with pre-warmed PBS and set on a temperature-controlled stage at 37 ℃. The force-distance curves were recorded and the elastic modulus of cells was calculated by NanoScope Analysis using the Hertz model.

### Western blotting

Equal amounts of total protein (50 μg) from each sample were separated in a 10% SDS-PAGE gel and transferred to a PVDF membrane at 120 V for 2 hours at room temperature. The blot was blocked with 5% nonfat dry milk suspended in 1x TBS (25 mM Tris, 137 mM NaCl, and 2.7 mM KCl) for 1 hour. Membranes were incubated sequentially with primary antibodies and secondary antibodies. Bands were scanned using a densitometer (Bio‐Rad) and quantified using the Quantity One 4.6.3 software (Bio‐Rad).

### Statistics

All data are presented as mean ± one standard deviation, where sample size (n) ≥ 3. Comparisons among values for groups greater than two were performed by using a one-way analysis of variance (ANOVA) followed by a Tukey’s post-hoc test. For two group analysis, a two-tailed, unpaired Student’s t-test was used to analyze differences. For all cases, p-values less than 0.05 were considered statistically significant. Origin 2018 software was used for all statistical evaluations.

## Data Availability

The authors declare that all data supporting the findings of this study are available within the paper and supplementary information files.

## Acknowledgements

We thank Dr. Marius Wernig at Stanford University for providing the constructs of BAM for reprogramming experiments, and thank Dr. Chih-Ming Ho for his suggestions on the design of microdevices. The authors were supported in part by a UCLA Eli and Edythe Broad Center of Regenerative Medicine and Stem Cell Research Innovation Award, a grant from the National Institute of Health (HL121450 to S.L. and GM140106 to S.K.), and the National Science Foundation (BMMB-1906165 to A.C.R.). The authors acknowledge the use of instruments at the Nano & Pico Characterization Lab and Advanced Light Microscopy and Spectroscopy Lab at the California NanoSystems Institute. The content is solely the responsibility of the authors and does not necessarily represent the official views of the National Institutes of Health.

## Author Contributions

Y.S., S.L., P.W. and S.K. designed the experiments. Y.S., J.S., B.C., W.K., N.Z., Q.P. and C.L. performed the experiments. Y.S., J.S., B.C., W.K., T.H., Q.P. and S.L. analyzed the data. Y.S., P.W., Y.W., A.C.R., C.H., S.K. and S.L. contributed to data interpretation and discussion. Y.S., J.S. and S.L. wrote the manuscript.

## Supplementary Information

Supplementary Figures and Table S1 can be found in Supplementary Information.

