## Supplementary Figures for "Transient nuclear deformation primes epigenetic state and promotes cell reprogramming"

a

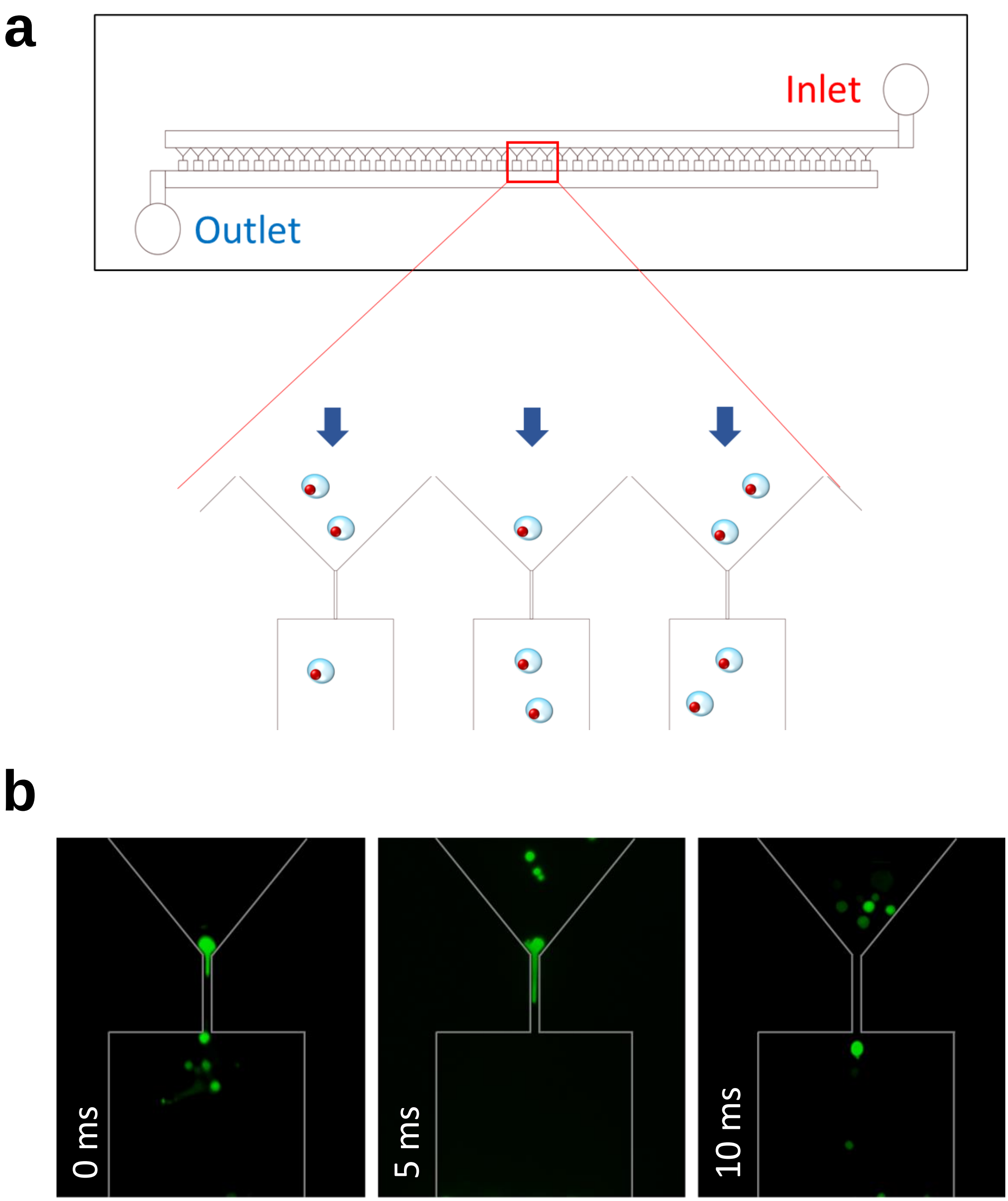


**Supplementary Figure S1. A microdevice with parallel microchannels to force transient cell deformation. (a)** Schematic illustrating the design of a microdevice with multiple microchannels to deform cells in parallel. **(b)** Fibroblasts were labeled with Cell Tracker Green, and live cell imaging was performed to monitor cells passing through the microchannels.

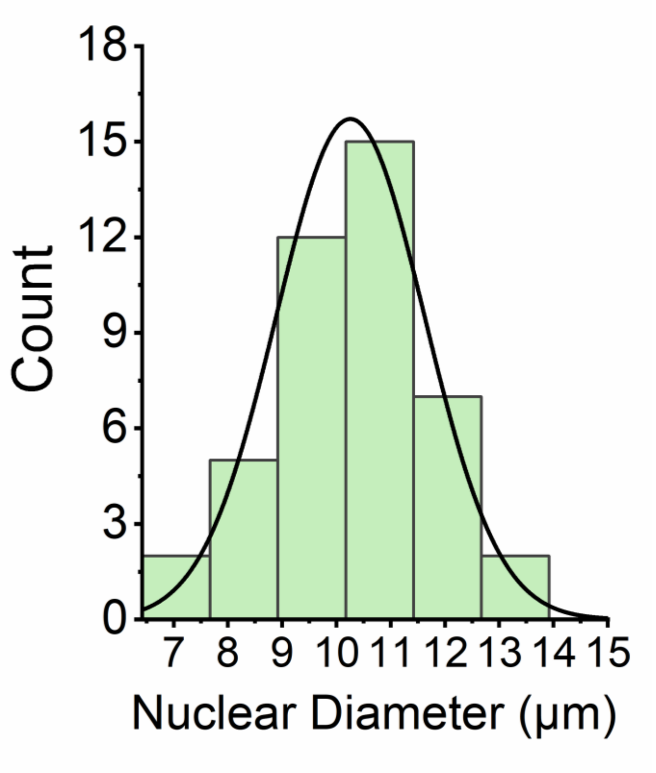


**Supplementary Figure S2**. **Nuclear diameter of cells passing 200-μm channel.** Living cells with nucleus staining (by Hoechst) were introduced into 200-μm channels. Epifluorescence images were collected by using a Zeiss Axio Observer Z1 inverted fluorescence microscope. Nuclear diameter was measured by the line tool of Zen Zeiss software (n=40).

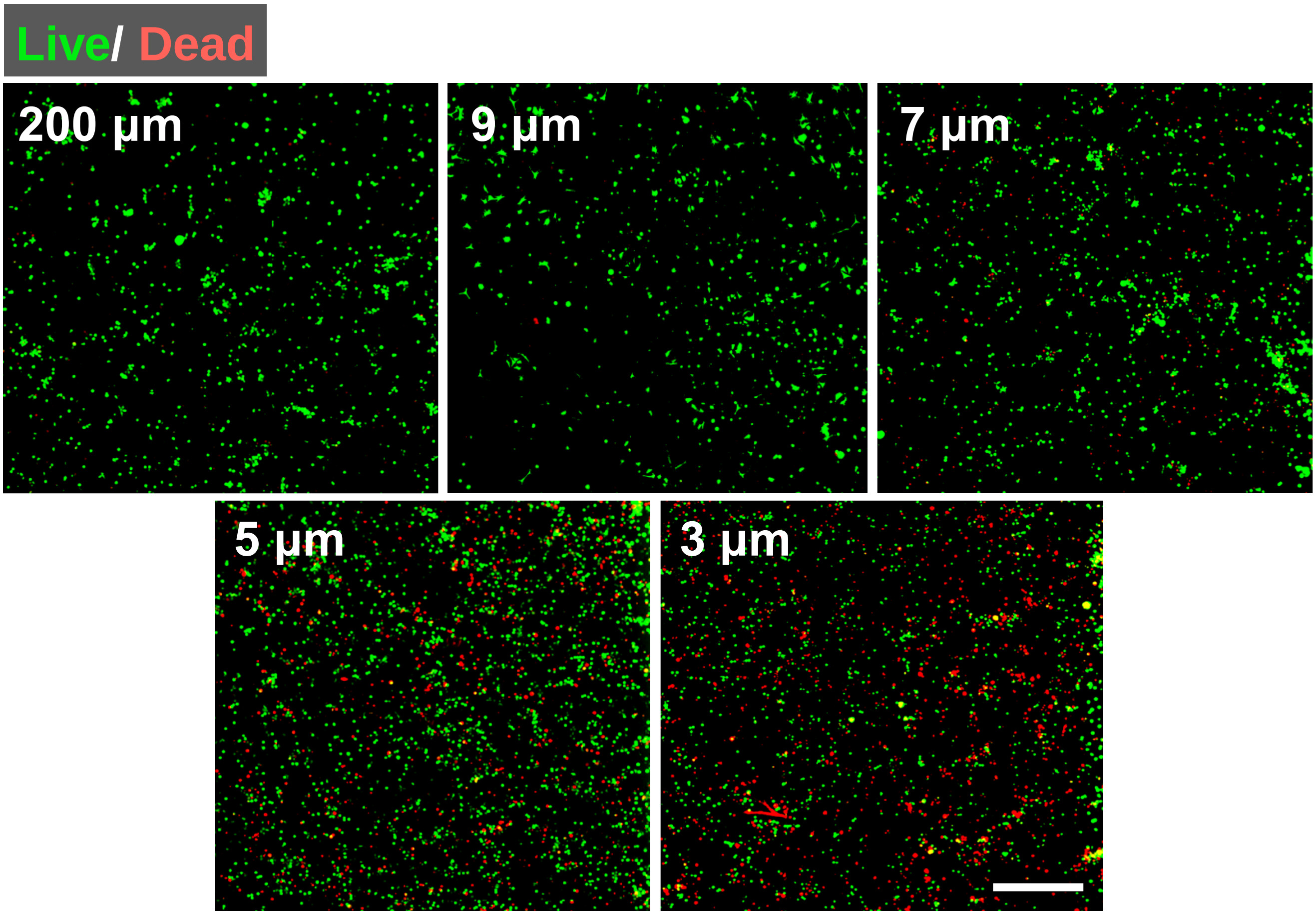


**Supplementary Figure S3. Effect of microchannel width on cell viability.** Immunofluorescent images show the viability of fibroblasts at 3 hours after passing through microchannels of various widths as determined by the LIVE/DEAD assay. Scale bar, 1 mm.

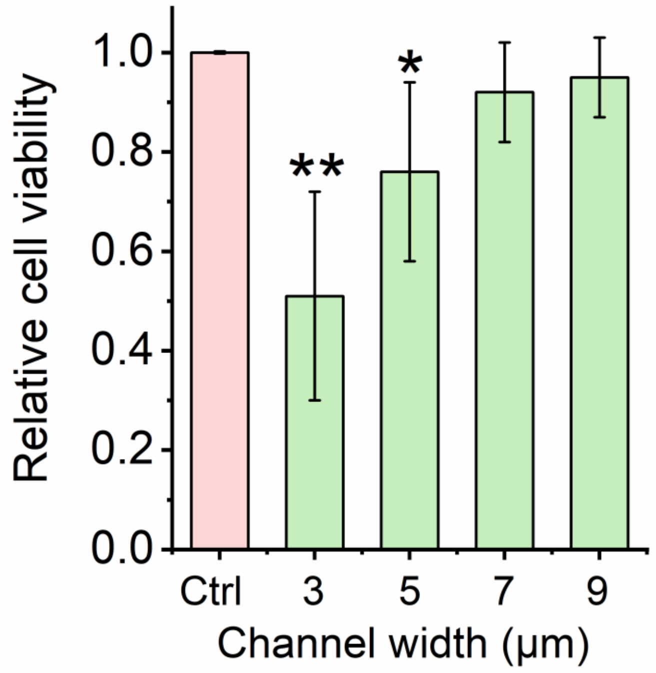


**Supplementary Figure S4. Effect of microchannel width on cell viability.** Cell viability of fibroblasts at 24 hours after passing through the microchannels of various widths as determined by the PrestoBlue® Cell Viability Reagent. Cell viability was normalized with the control (The viability of cells passing through 200-μm channels). Bar graph shows mean ± SD (n=6, *p<0.05, **p<0.01, compared with control). Statistical significance was determined by a one-way ANOVA and Tukey's multiple comparison test.

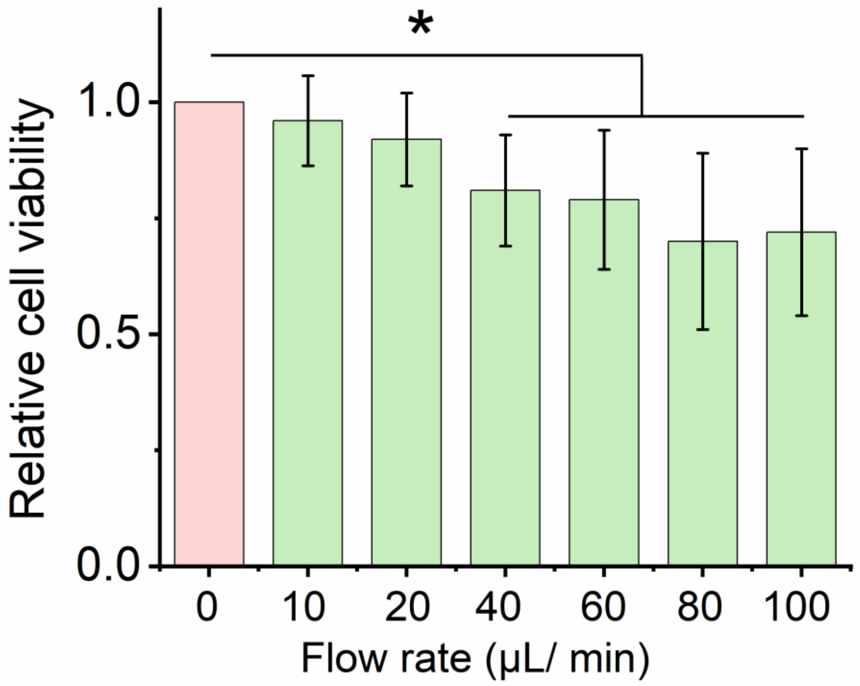


**Supplementary Figure S5. Effect of flow rate on cell viability.** Quantification of cell viability at distinct flow rates (normalized to the 0 μL/min flow rate group). Cells cultured under static condition were set as the 0 μL/minute flow rate group. Bar graphs show mean ± standard deviation (n=6; *$p<0.05$ in comparison with the 0 μL/minute flow rate group). Statistical significance was determined by a one-way ANOVA and Tukey's multiple comparison test.

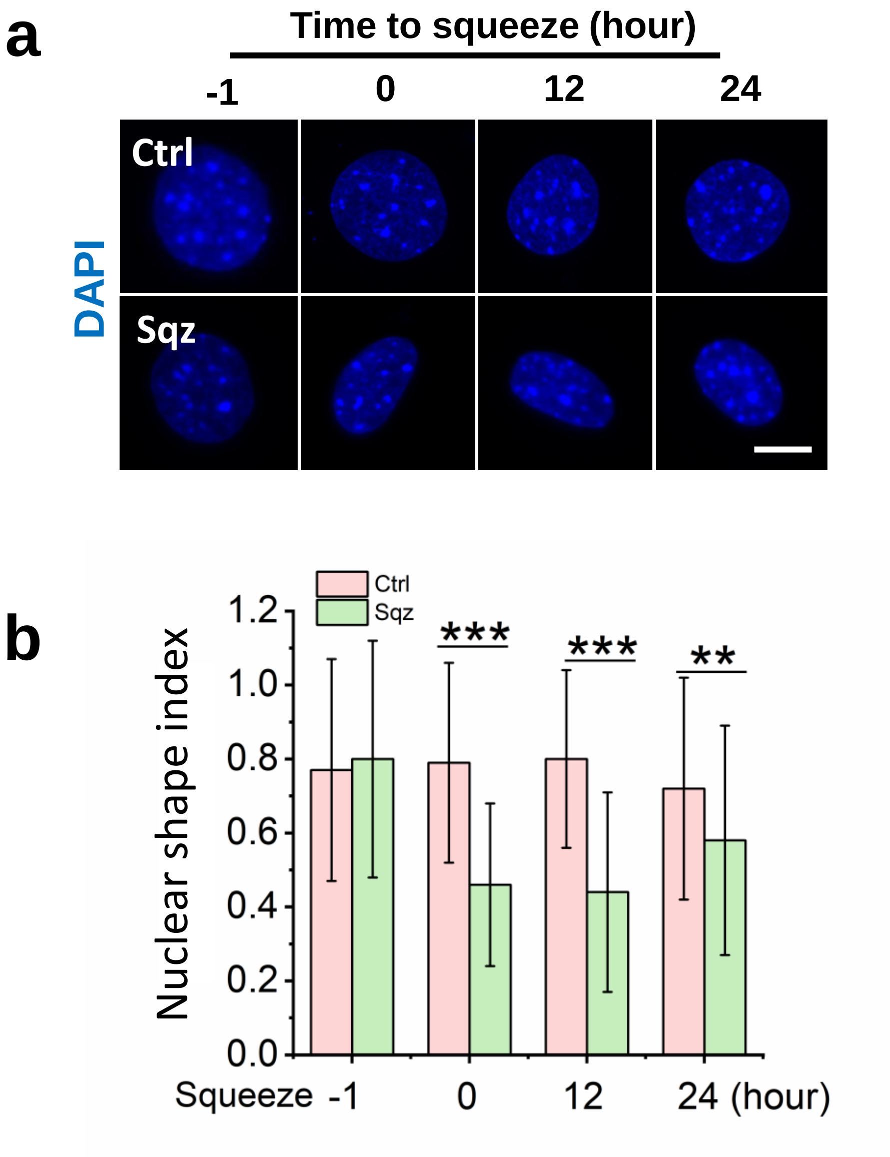


**Supplementary Figure S6. Microchannel-induced nuclear morphology changes. (a)** Immunofluorescent images of nuclei before and after cells were squeezed through the microchannel. Control (Ctrl): cells passing through 200-μm channels. Squeezed (Sqz): cells passing through 7-μm microchannels. Scale bar, 10 μm. **(b)** Quantification of nuclear shape index in control and mechanically squeezed cells at the indicated time points. Nuclear perimeter (C) and projected area (A) were assessed using Image J software based on the fluorescent images. Nuclear shape index is defined as $4\pi \times A/C^2$. Bar graph shows mean ± SD ($n \geq 50$, $^{**}p<0.01$, $^{***}p<0.001$). Statistical significance was determined at each time point by a two-tailed, unpaired t-test.

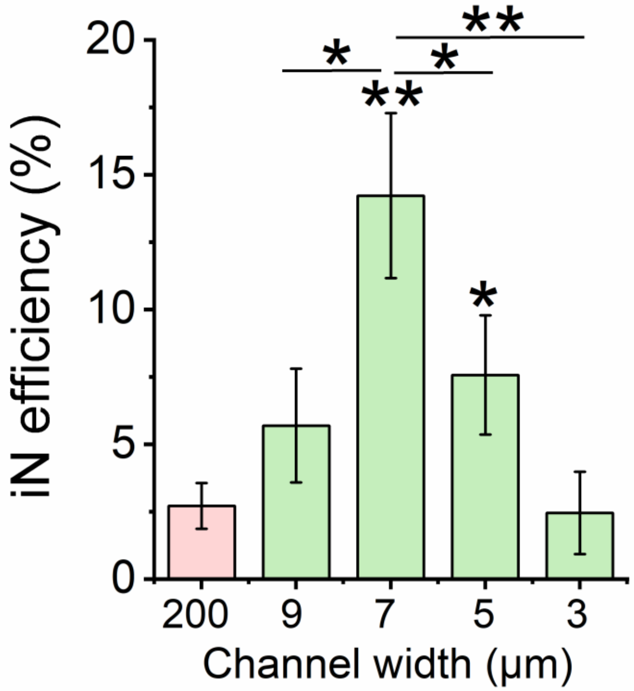


**Supplementary Figure S7. Effect of microchannel width on iN reprogramming efficiency.** Reprogramming efficiency of BAM-transduced fibroblasts at the day 7 after passing through the microchannels of various widths was determined by the Tuj1 staining. Cells pass through the 200-μm channels were used as a control. Bar graph shows mean ± SD (n=3, *p<0.05, **p<0.01 compared with the control). Statistical significance was determined by a one-way ANOVA and Tukey's multiple comparison test.

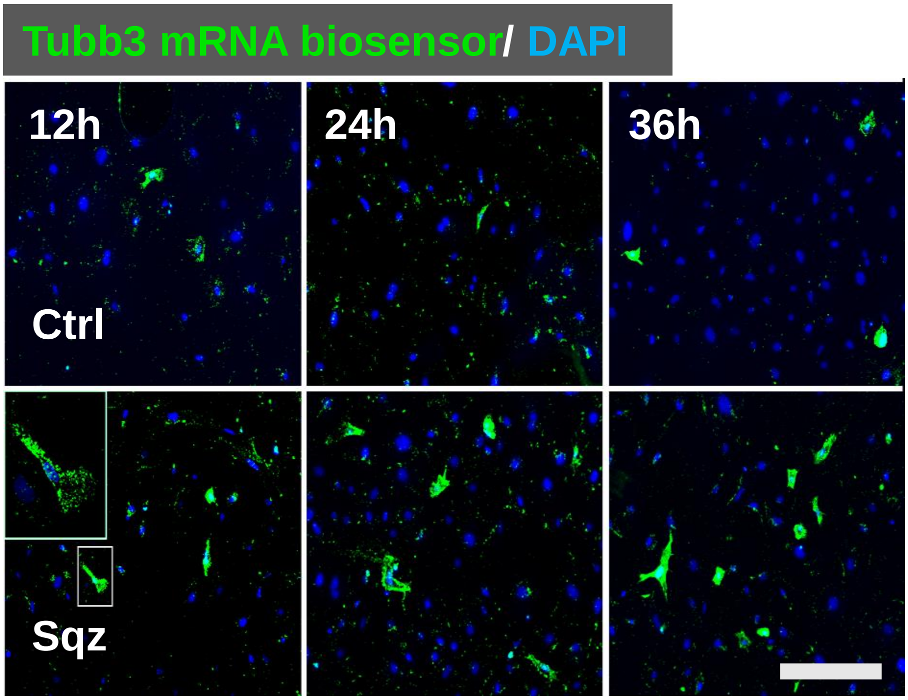


**Supplementary Figure S8. Nuclear deformation promoted Tubb3 gene expression.** At 12, 24 and 36 hours after BAM-transduced fibroblasts passed through the microchannels, Tubb3 mRNA biosensor were added into culture medium and incubated for 4 hours before taking images. The samples were washed by PBS 3 times, and images were taken by immunofluorescence microscopy. Immunofluorescent images show Tuj1 mRNA expression in the cells of the control (Ctrl) and squeezed (Sqz) groups at the indicated time points after mechanical squeezing, as detected by a Tubb3 mRNA biosensor and immunofluorescence microscopy. Scale bar, 100 µm.

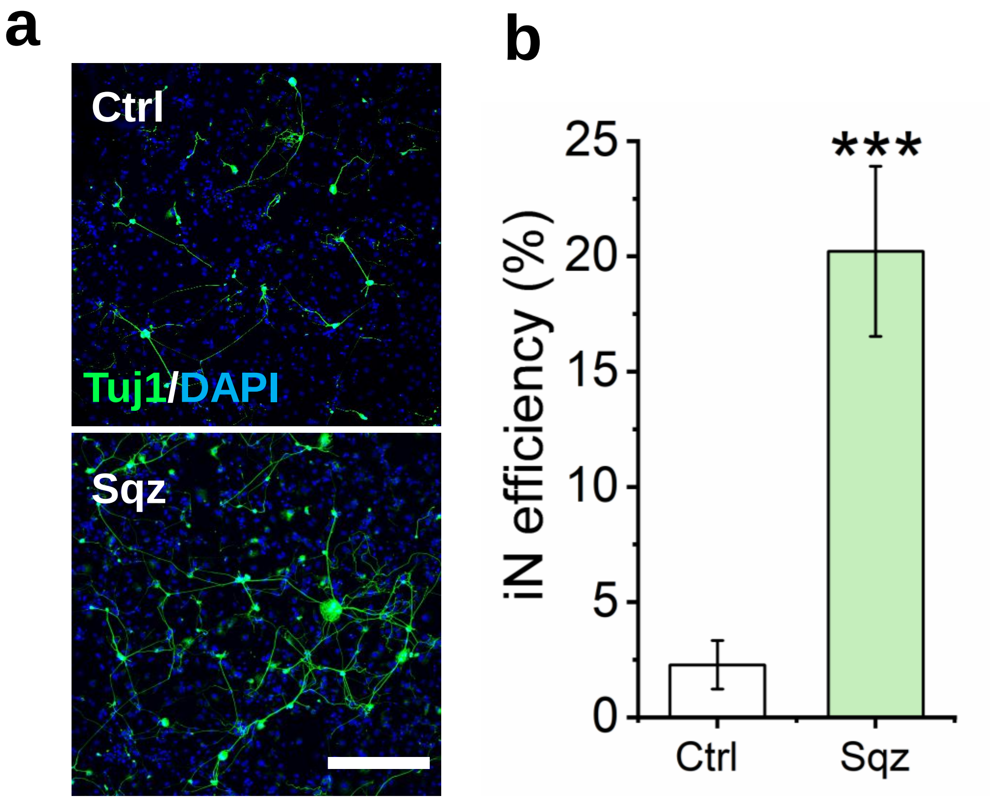


**Supplementary Figure S9. Nuclear deformation promoted Tuj1 expression. (a)** Immunofluorescent images of Tuj1+ iN cells on day 14. Scale bar, 200 µm. **(b)** Reprogramming efficiency of BAM-transduced fibroblasts in the control (Ctrl) and squeezed (Sqz) groups at day 14. Bar graphs show mean ± standard deviation (n=6; ***$p<0.001$); significance was determined by a two-tailed, unpaired t-test.

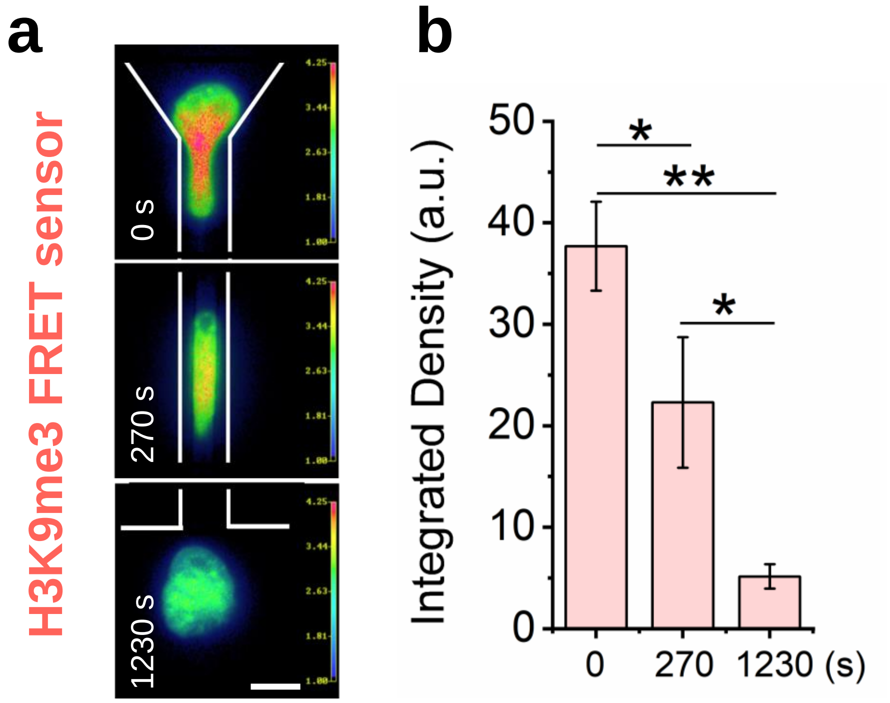


**Supplementary Figure S10. Cell nucleus deformation in microchannel decreased H3K9me3 as detected by a FRET sensor. (a)** Fibroblasts were transduced with viruses to express a H3K9me3 FRET sensor. Live cell imaging was performed to monitor changes in H3K9me3 during cell deformation induced by the microchannel. Red color indicates higher level of H3K9me3, and green indicates lower FRET signal. **(b)** H3K9me3 FRET signal was quantified at different time point. Bar graph shows mean ± standard deviation (n=3; **p<0.01); statistical significance was determined by a one-way ANOVA and Tukey's multiple comparison test.

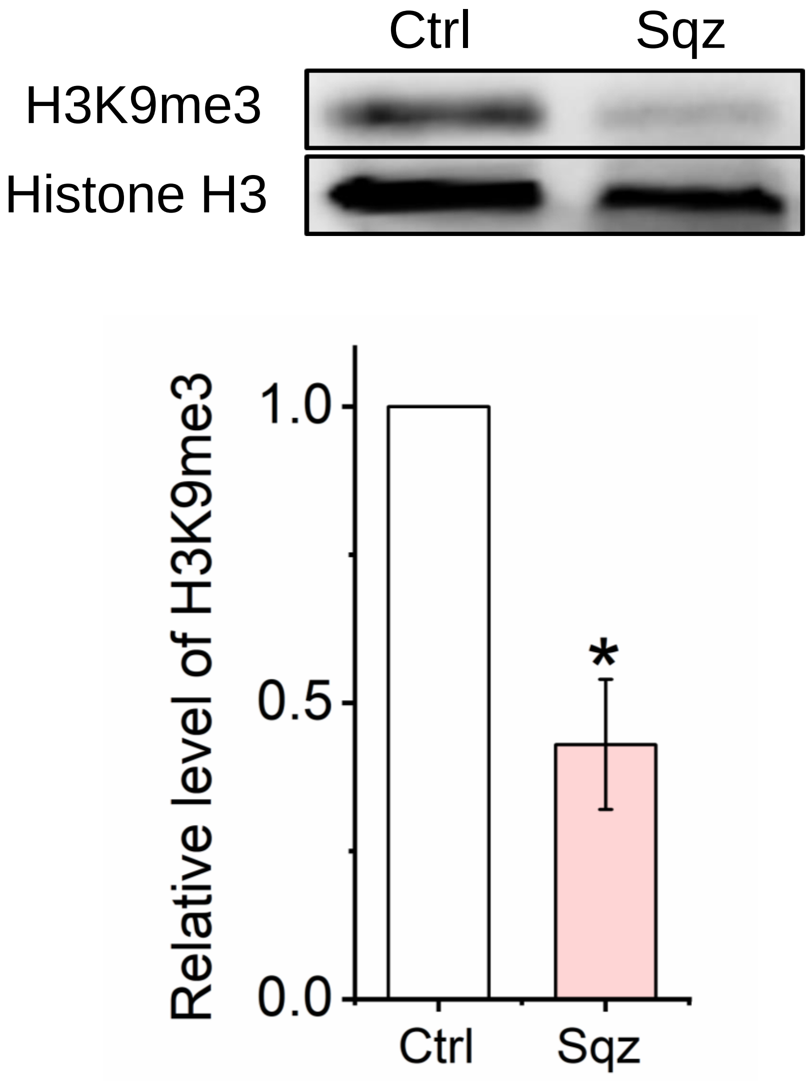


**Supplementary Figure S11. Microchannel-induced nuclear deformation decreased global H3K9me3.** (a) Total histone extracts were isolated from BAM-transduced fibroblasts by Histone Extraction Kit (Abcam, ab221031, USA) at 3 hours after the cells passed through the microchannels. Western blotting was used to examine the levels of H3K9me3 and histone H3. Control (Ctrl): cells passing through 200-μm channels. Squeezed (Sqz): cells passing through 7-μm microchannels. (b) Quantification of H3K9me3 from blot. Bar graph shows mean ± standard deviation (n=3; *p<0.05); statistical significance was determined by a one-way ANOVA and Tukey's multiple comparison test.

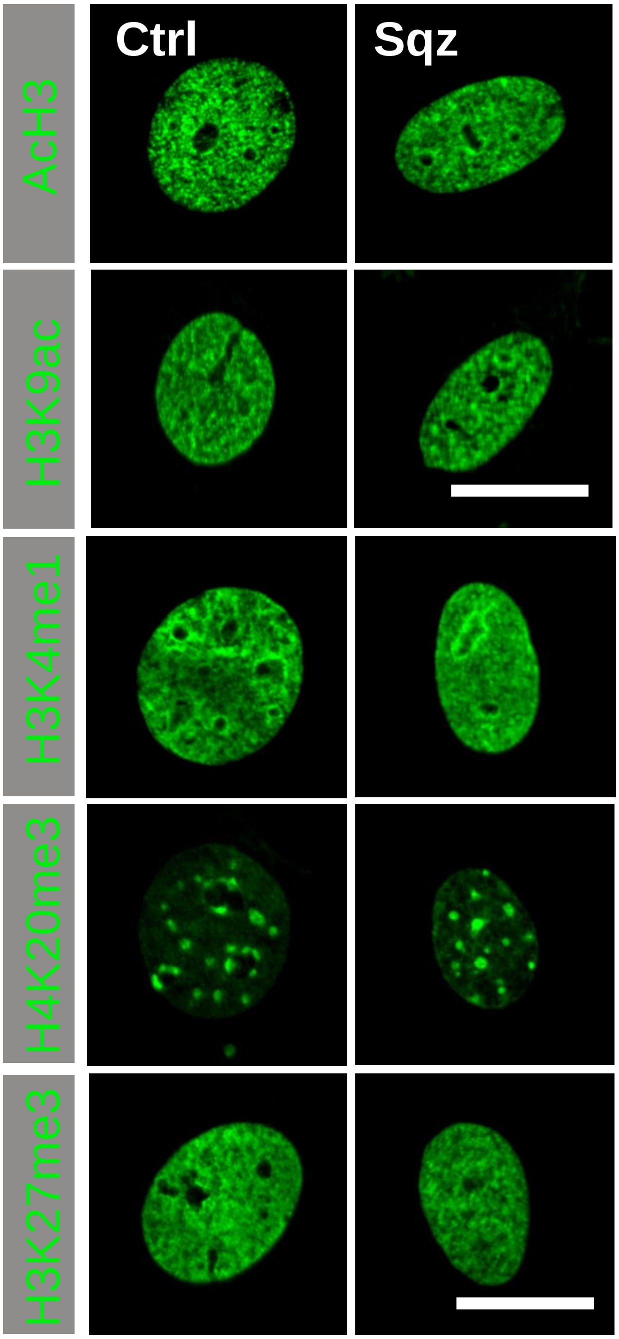


**Supplementary Figure S12. Effect of forced nuclear deformation on histone marks.** Representative immunofluorescent images show the level and distribution of various histone marks in BAM-transduced fibroblasts 3 hours after the cells passed through the microchannels. Control (Ctrl): cells passing through 200-μm channels. Squeezed (Sqz): cells passing through 7-μm microchannels. Scale bar, 10 μm.

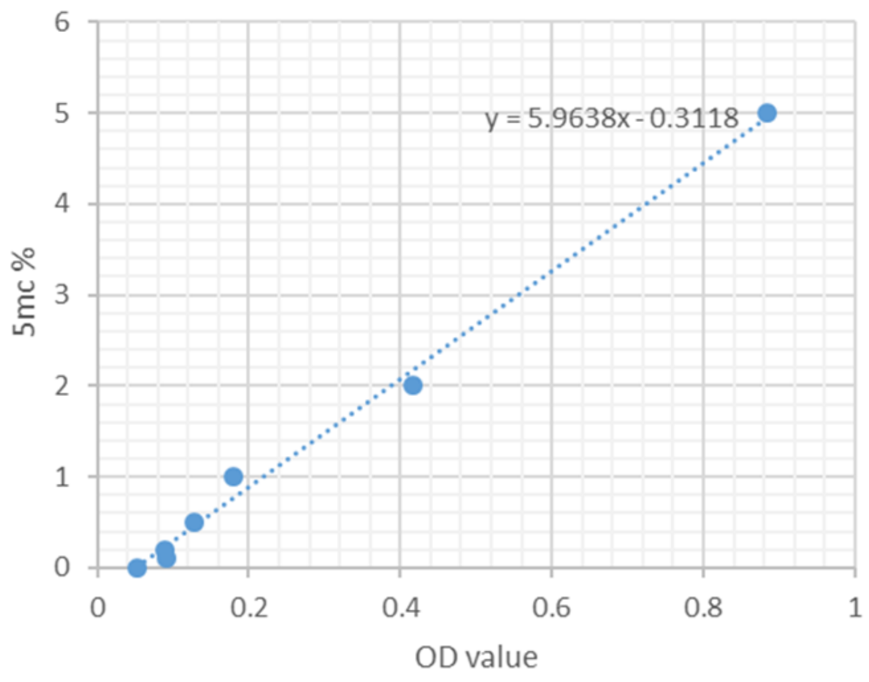


**Supplementary Figure S13. 5-mC standard curve.** The 5-mC standard sample was diluted to 0.1% to 5% (1 μg/ml to 50 μg/ml) according to the manufacturer's instructions for the MethylFlashTM Global DNA Methylation (5-mC) ELISA Easy Kit (Epigentek, P-1030), and bonded into the assay wells and incubated with 5-mC detection complex solution for 60 minutes. The color developer solution was added into assay wells, and absorbance was detected by using a plate reader (Inifinte 200Pro, 30050303) at 450 nm. Based on the average OD value on the X-axis and the known 5-mC percentage at each point on the Y-axis, we generated the standard curve between 5-mC % and OD value. This standard curve was used to calculate the 5-mC level in cells.

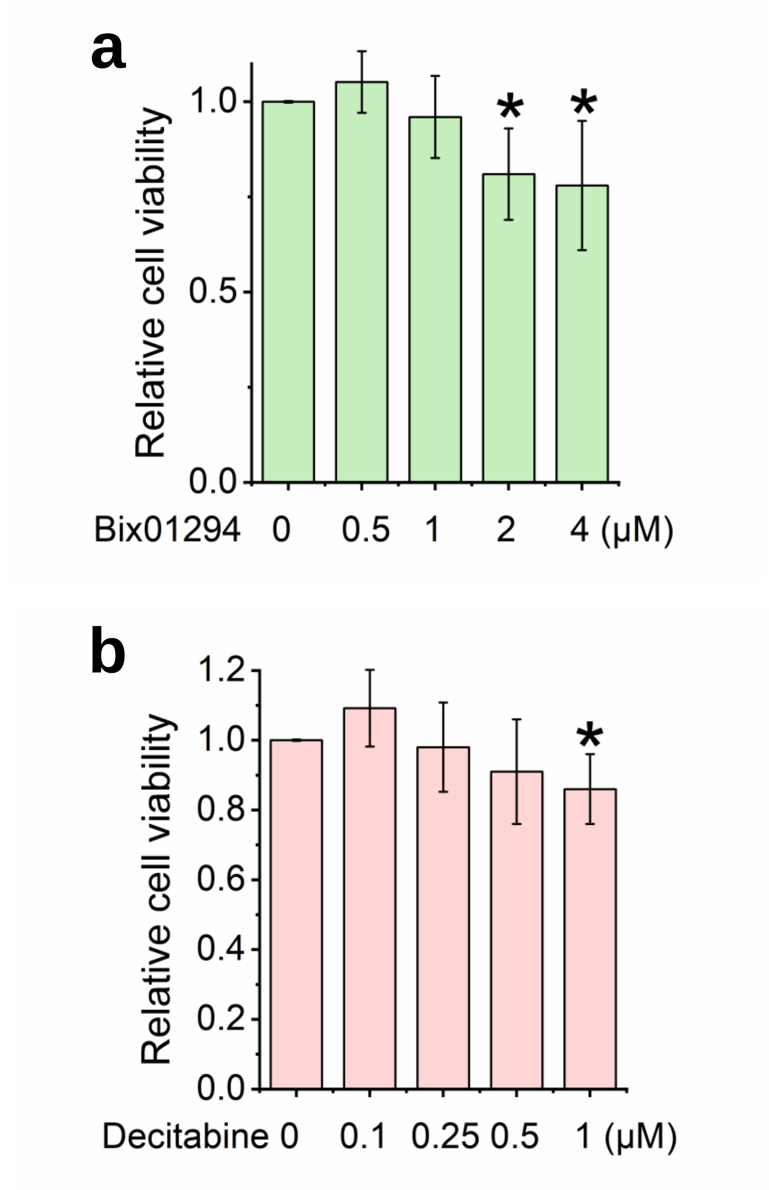


**Supplementary Figure S14. Effects of histone methyltransferase (HMT) and DNA methyltransferase (DNMT) inhibition on cell viability.** (**a**) Fibroblasts were treated with HMT inhibitor Bix01294 at various concentrations for 24 hours, and cell viability was determined by using the PrestoBlue® Cell Viability Reagent. (**b**) Fibroblasts were treated with DNMT inhibitor Decitabine at various concentrations for 24 hours, followed by cell viability assay. Bar graph shows mean ± SD (n=3, *$p<0.05$ compared to solvent control without inhibitor). Cell viability was normalized with the solvent control (the viability of cells treated by DMSO). Statistical significance was determined by a one-way ANOVA and Tukey's multiple comparison test.

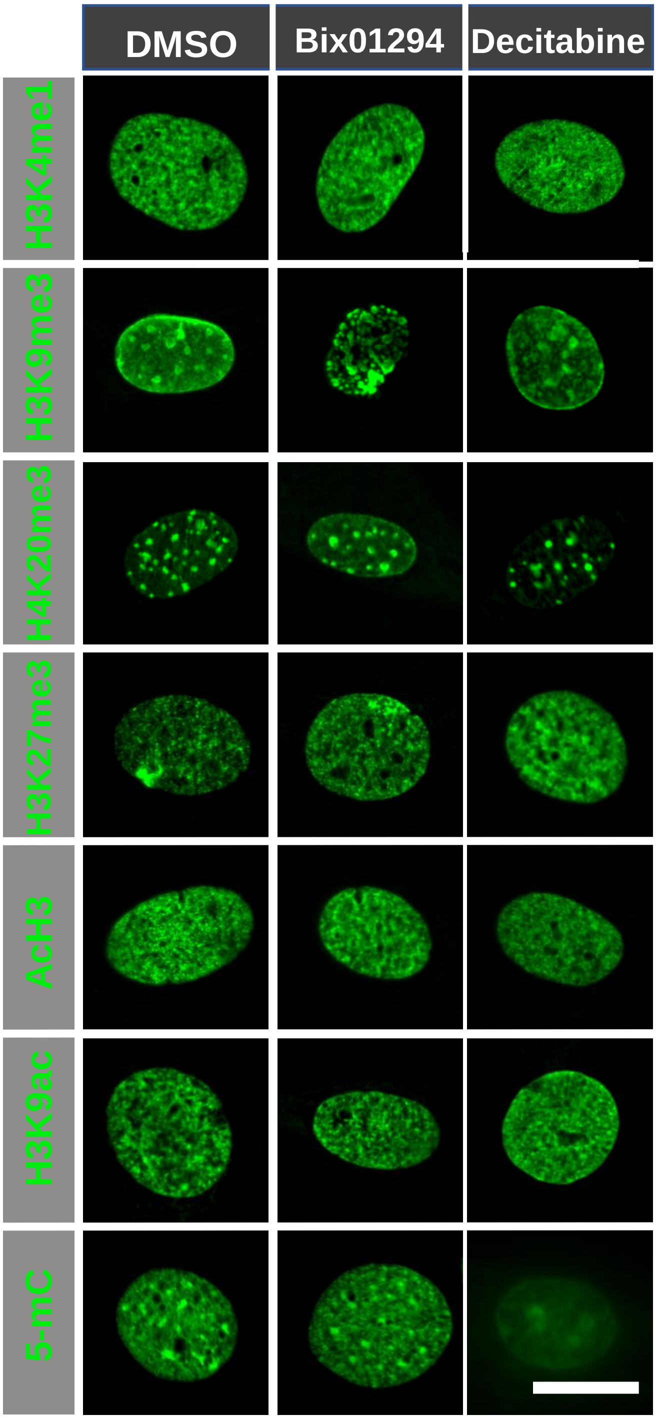


**Supplementary Figure S15. Effect of HMT and DNMT inhibition on histone marks.** Fibroblasts were treated with DMSO (solvent control), H3K9me3-specific HMT inhibitor Bix01294, or DNMT inhibitor Decitabine for 24 hours. Cells were fixed and stained for various histone marks. Representative immunofluorescent images were collected by fluorescence microscopy. Scale bar, 10 µm.

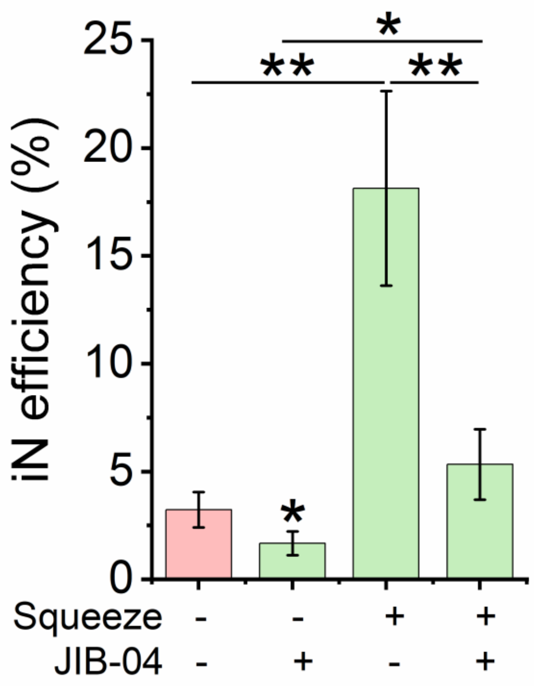


**Supplementary Figure S16. Effect of HDMT inhibition on iN reprogramming efficiency.** BAM-transduced fibroblasts were pretreated with JIB-04 (100 nM) for 24 hours before passing through the microchannels. Reprogramming efficiency was determined by the Tuj1 staining. Cells passing through the 200-μm channels were used as a control. Bar graph shows mean ± SD (n=3, *p<0.05, **p<0.01 compared with the control). Statistical significance was determined a one-way ANOVA and Tukey's multiple comparison test.

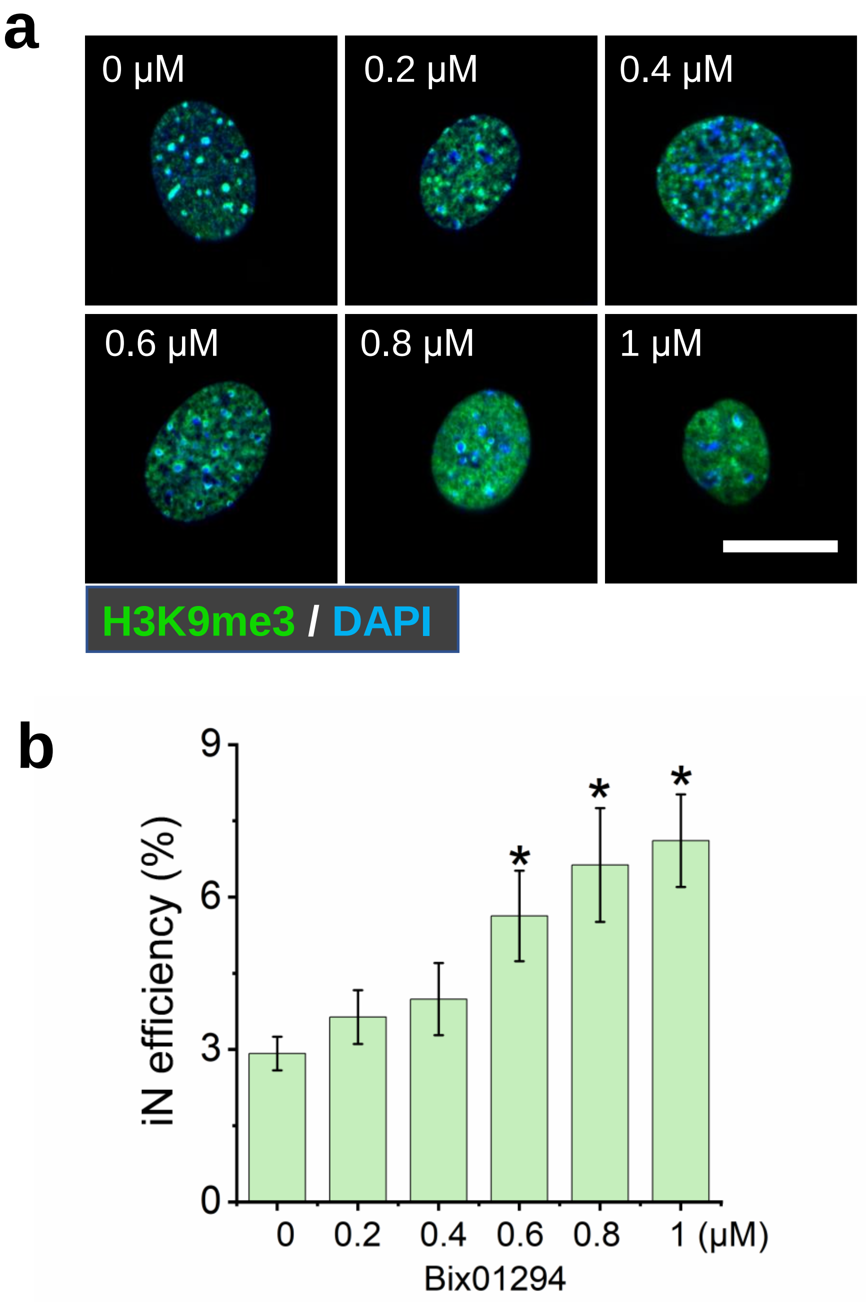


**Supplementary Figure S17. Effect of HMT inhibition on histone marks and iN reprogramming efficiency. (a)** Representative immunofluorescent images show the level and distribution of H3K9me3 in BAM-transduced fibroblasts pre-treated with different doses of HMT inhibitor (Bix01294) for 24 hours. Bix01294 specifically inhibits H3K9me3. Scale bar, 10 μm. **(b)** Reprogramming efficiency of BAM-transduced fibroblasts that were pretreated with Bix01294 (0.2-1 μM) for 24 hours before adding Dox. Cells treated by DMSO were used as a control. Bar graph shows mean ± SD (n=3, **p<0.01 compared with the control). Statistical significance was determined by a one-way ANOVA and Tukey's multiple comparison test.

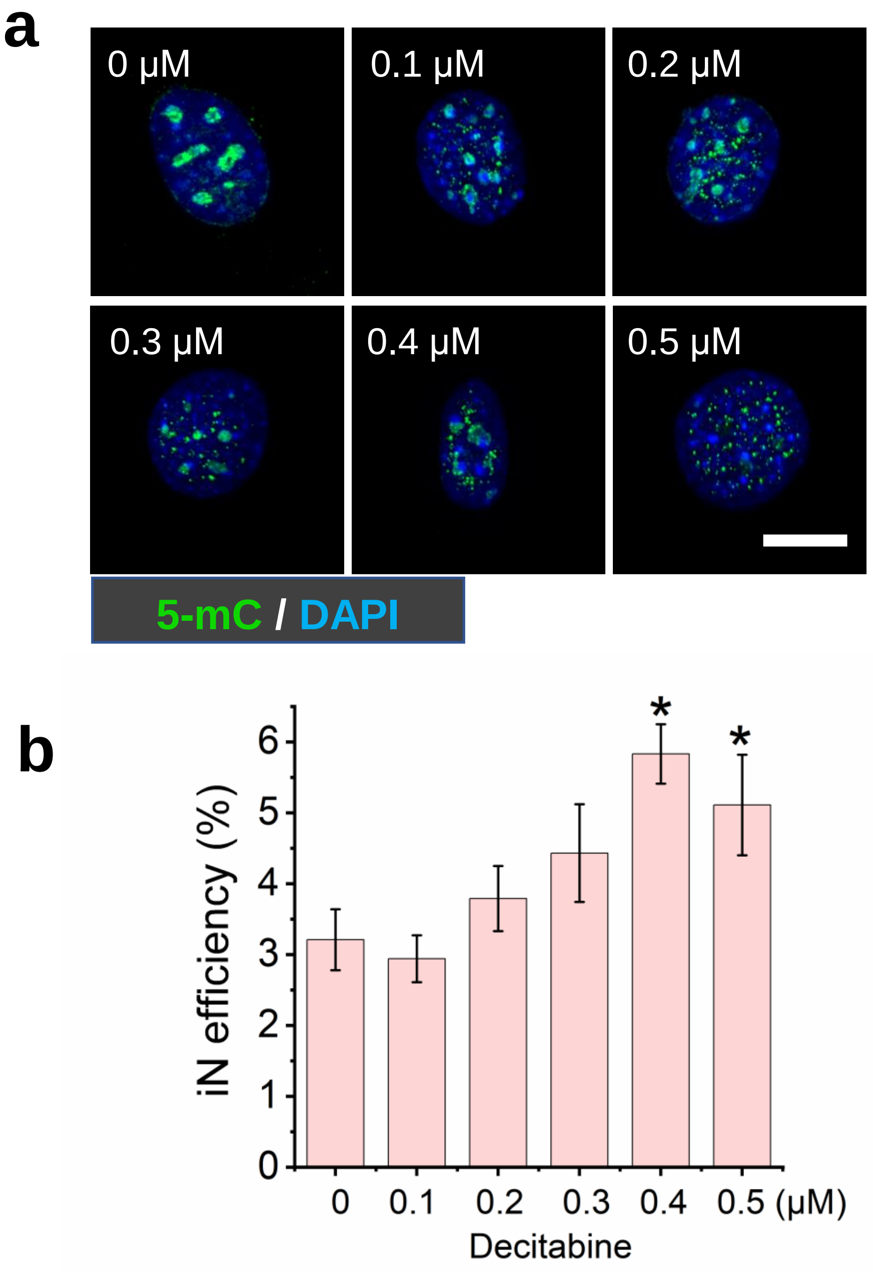


**Supplementary Figure S18. Effect of DNMT inhibition on histone marks and iN reprogramming efficiency. (a)** Representative immunofluorescent images show the level and distribution of 5-mC in BAM-transduced fibroblasts pre-treated with different doses of DNMT inhibitor (Decitabine) for 24 hours. Decitabine specifically inhibits 5-mC. Scale bar, 10 μm. **(b)** Reprogramming efficiency of BAM-transduced fibroblasts that were pretreated with Decitabine (0.1-0.5 μM) for 24 hours before adding Dox. Cells treated by DMSO were used as a control. Bar graph shows mean ± SD (n=3, $^{**}p<0.01$ compared with the control). Statistical significance was determined by a one-way ANOVA and Tukey's multiple comparison test.

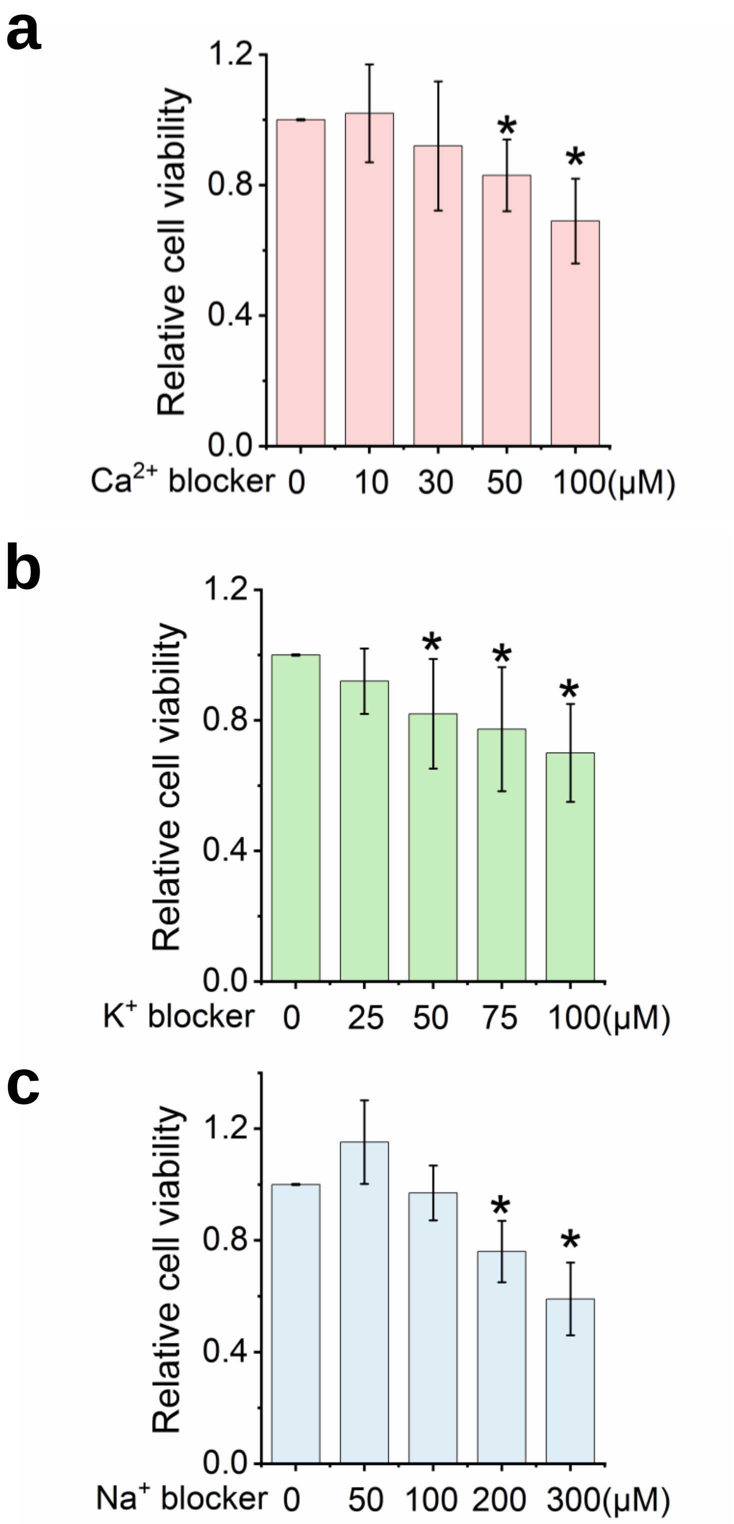


**Supplementary Figure S19. Effect of ion channel inhibition on cell viability.** Fibroblasts were treated with blockers of calcium (**a**), potassium (**b**), and sodium (**c**) ion channels, respectively, at the indicated concentrations for 12 hours before adding Dox and passing the cells through the microchannel as determined by the PrestoBlue® Cell Viability Reagent. Cells treated by DMSO were used as control. Bar graph shows mean ± SD (n=3, *p<0.05 compared with the control). Statistical significance was determined by a one-way ANOVA and Tukey's multiple comparison test.

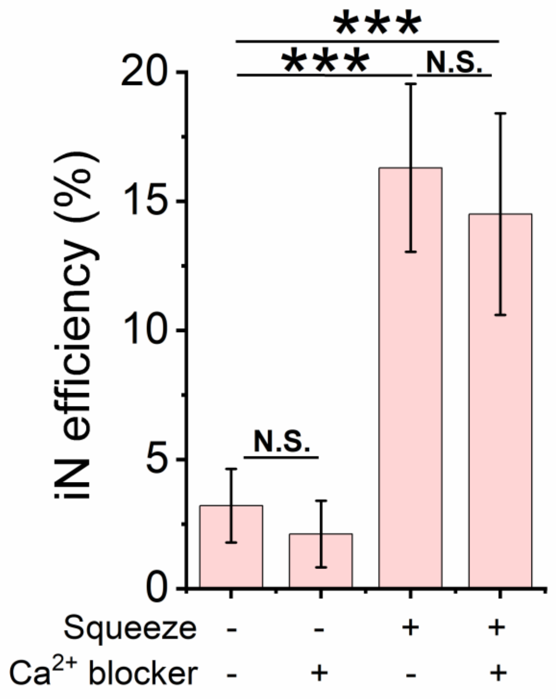


**Supplementary Figure S20. Calcium ion channels did not regulate microchannel-induced iN reprograming.** Reprogramming efficiency of BAM-transduced fibroblasts that were pretreated with the calcium channel blocker Amlodipine (30 μM ) for 12 hours before adding Dox and squeezing the cells with the microchannel. Bar graph shows mean ± SD (n=6, ***p<0.001). Statistical significance was determined by a one-way ANOVA and Tukey's multiple comparison test.

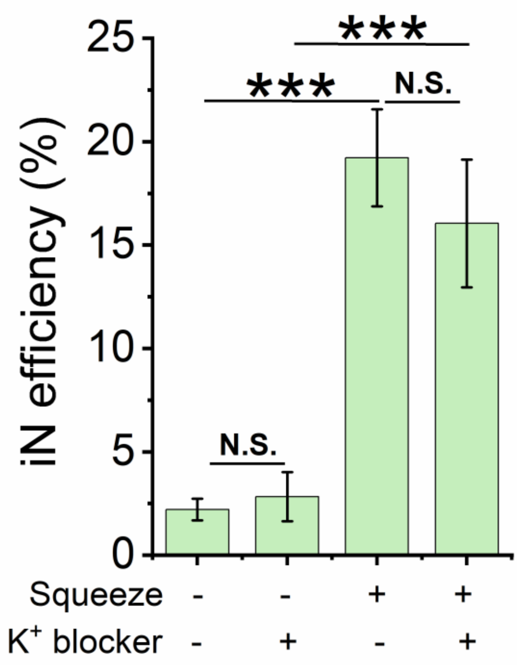


**Supplementary Figure S21. Potassium ion channels did not significantly affect microchannel-induced iN reprograming.** Reprogramming efficiency of BAM-transduced fibroblasts that were pretreated with the potassium channel blocker Quinine (25 μM) for 12 hours before adding Dox and passing the cells through the microchannels. Bar graph shows mean ± SD (n=6, ***$p<0.001$). Statistical significance was determined by a one-way ANOVA and Tukey's multiple comparison test.

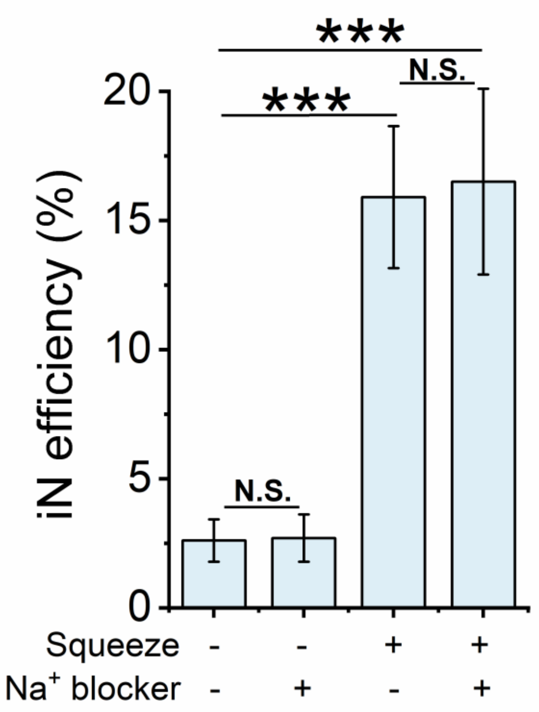


**Supplementary Figure S22. Sodium ion channels did not significantly affect microchannel-induced iN reprograming.** Reprogramming efficiency of BAM-transduced fibroblasts that were pretreated with the sodium channel blocker procainamide (100 μM) for 12 hours before adding Dox and passing the cells through the microchannels. Bar graph shows mean ± SD (n=6, ***p<0.001). Statistical significance was determined by a one-way ANOVA and Tukey's multiple comparison test.

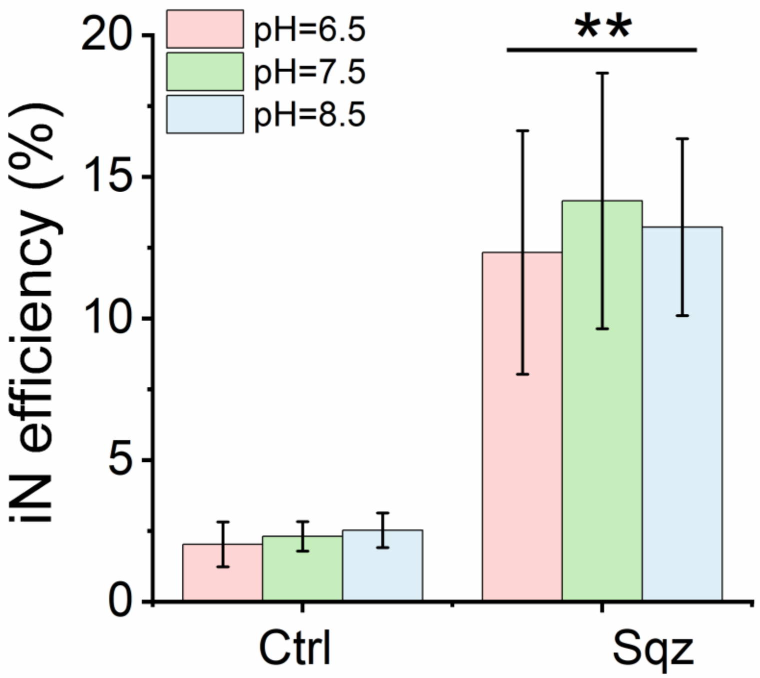


**Supplementary Figure S23. Extracellular pH did not affect microchannel-induced iN reprograming.** Reprogramming efficiency of BAM-transduced fibroblasts that were pretreated with culture media at different pH for 1 hour before the mechanical deformation induced by the microchannels. Control (Ctrl): cells passing through 200-μm channels. Squeezed (Sqz): cells passing through 7-μm microchannels. Bar graph shows mean ± SD (n=3, *p<0.05 compared with the control). Statistical significance was determined by a one-way ANOVA and Tukey's multiple comparison test.

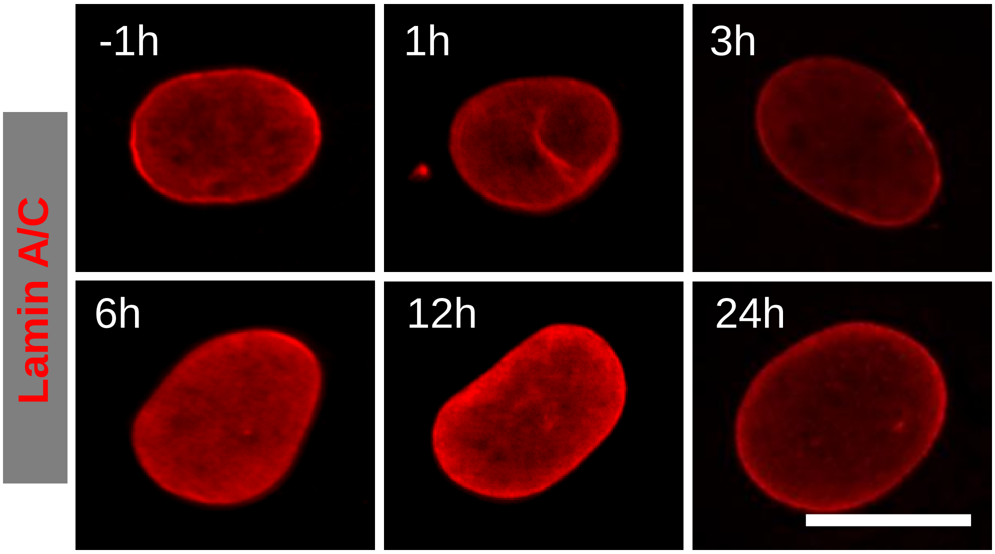


**Supplementary Figure S24. Characterization of lamin A/C before and after mechanical deformation.** Representative images of lamin A/C staining in cells at the indicated time points before or after passing through 200-μm channels. Scale bar, 10 μm.

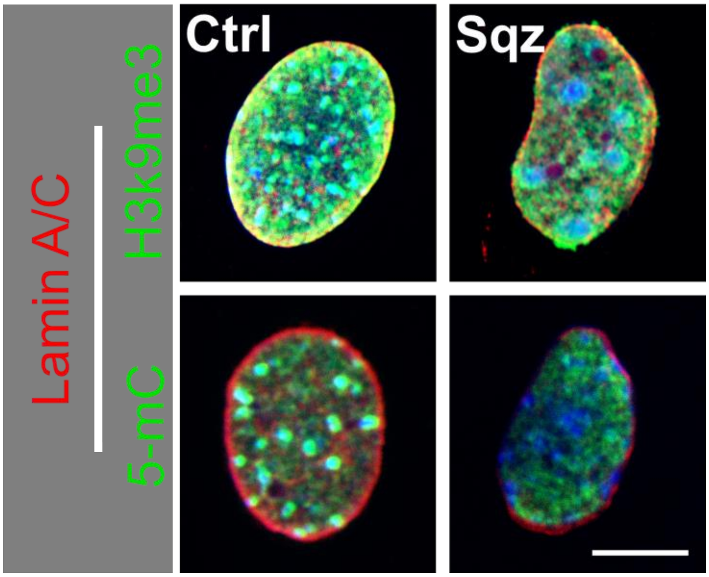


**Supplementary Figure S25. Characterization of lamin A/C, H3K9me3 and 5mC at 1 hour after the cells passed through microchannels.** Representative images of lamin A/C **,** staining in cells at the indicated time points after passing through 7-μm microchannels. Control (Ctrl): cells passing through 200-μm channels. Squeezed (Sqz): cells passing through 7-μm microchannels. Scale bar, 10 μm.

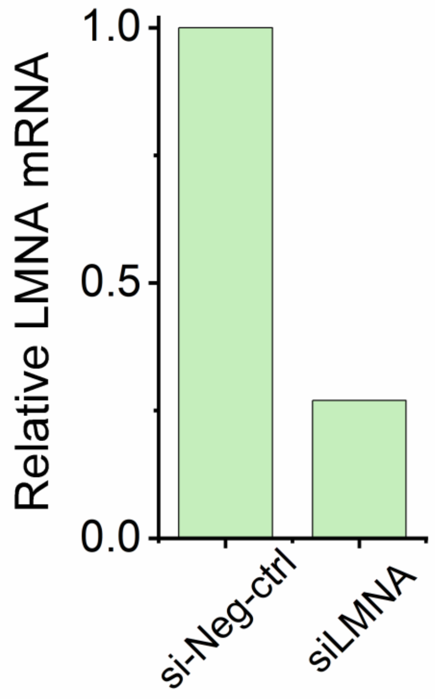


**Supplementary Figure S26. Lamin A knockdown by using siRNA interference.** Fibroblasts were transfected with an siRNA against lamin A (*LMNA*) or negative control, respectively for 12 hours, and RNA was isolated from the samples after 1 day. qRT-PCR analysis confirmed the successful knockdown of lamin A gene expression. Gene expression was normalized to 18S.

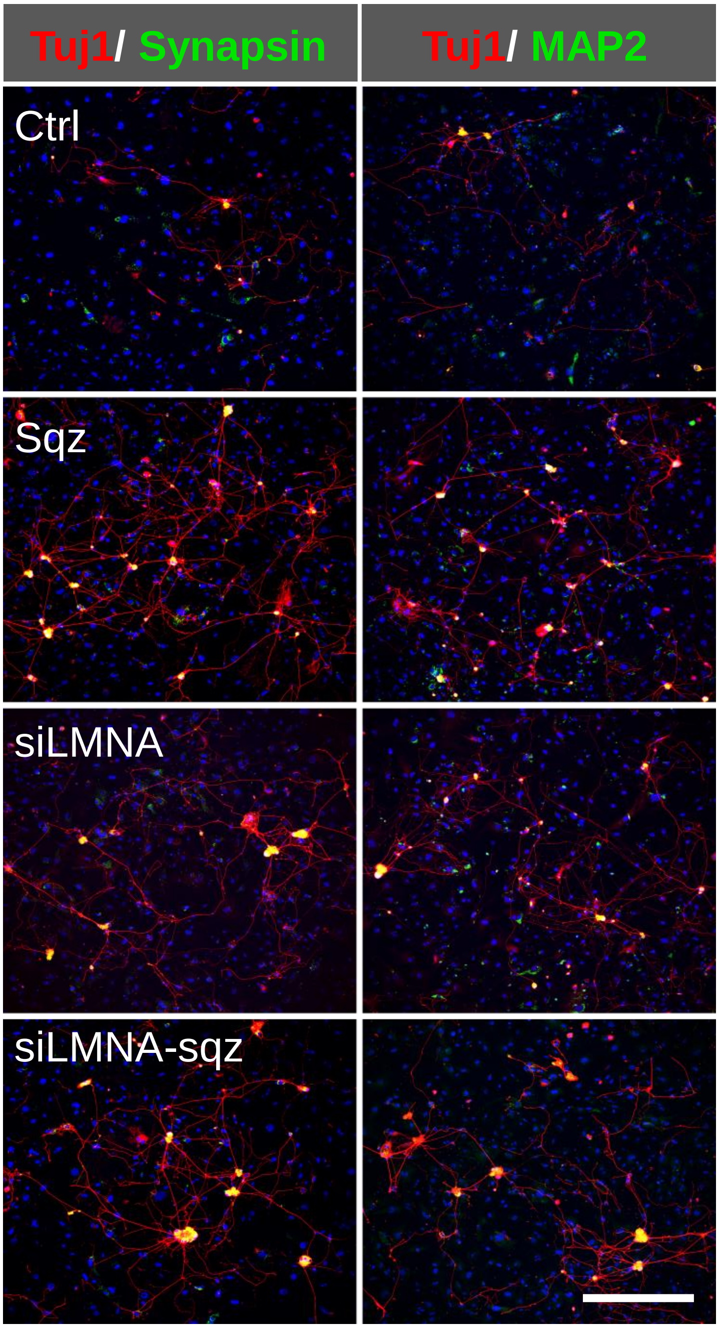


**Supplementary Figure S27. Lamin A regulated microchannel-induced iN reprogramming.** BAM-transduced fibroblasts were transfected with lamin A siRNA for 12 hours. Dox was added to culture media for 6 hours, and cells were detached and introduced into microchannels for nuclear deformation. Cells passing through 200-μm channels were used as a control. Representative images of Tuj1 (Red), MAP2 (Green) and synapsin (Green) staining in control and lamin A-knockdown fibroblasts at 4 weeks after nuclear deformation. DNA was stained by DAPI (Blue). Scale bar, 200 μm.

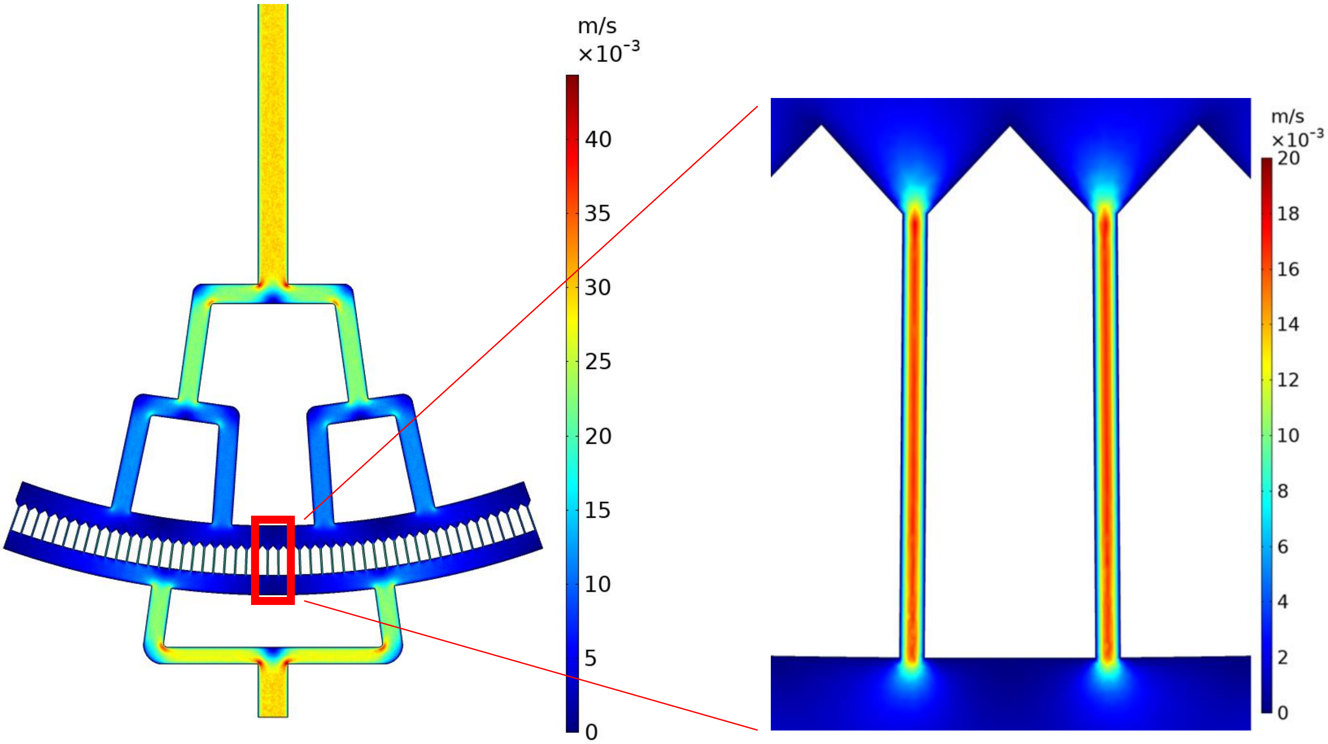


**Supplementary Figure S28.** Simulation of velocity magnitude in the high-throughput microfluidic device.

**Table S1. Antibodies used in immunocytochemistry**.

| Antibody | Vendor | Catalog # | Dilution |
|---|---|---|---|
| Tuj1 | Biolegend | 801202 | 1:1000 |
| H3K4me1 | Abcam | ab32356 | 1:300 |
| H3K9me3 | Abcam | ab8898 | 1:500 |
| H4K20me3 | Abcam | ab4729 | 1:300 |
| H3K27me3 | Abcam | ab192985 | 1:300 |
| AcH3 | Millipore | 06-599 | 1:300 |
| H3K9ac | Abcam | ab4441 | 1:300 |
| 5-mC | Millipore | NA81 | 1:300 |
| Synapsin | Abcam | ab64581 | 1:100 |
| MAP-2 | Sigma | M9942 | 1:200 |
| Lamin A/C | Santa Cruz | sc-376248 | 1:300 |
| Histone H3 | Santa Cruz | sc-8654 | 1:1000 |
